# F-actin regulates polarized secretion of pollen tube attractants in *Arabidopsis* synergid cell

**DOI:** 10.1101/2022.06.14.496136

**Authors:** Daichi Susaki, Rie Izumi, Takao Oi, Hidenori Takeuchi, Ji Min Shin, Naoya Sugi, Tetsu Kinoshita, Tetsuya Higashiyama, Tomokazu Kawashima, Daisuke Maruyama

**Author notes:** Footnotes: The author responsible for distribution of materials integral to the findings presented in this article in accordance with the policy described in the Instructions for Authors (https://academic.oup.com/plcell/pages/General-Instructions) is: Daisuke Maruyama.”.

## Abstract

Pollen tube attraction is a key event of sexual reproduction in flowering plants. In the ovule, two synergid cells neighboring the egg cell control pollen tube arrival via the active secretion of attractant peptides such as AtLURE1 and XIUQIU from the filiform apparatus facing toward the micropyle. Distinctive cell polarity together with longitudinal F-actin and microtubules are hallmarks of the synergid cell in various species, though functions of these cellular structures are still unclear. In this study we used genetic and pharmacological approaches to elucidate the roles of cytoskeletal components in filiform apparatus formation and pollen tube guidance in *Arabidopsis thaliana*. Inhibition of microtubule formation reduced invaginations of the plasma membrane but did not abolish micropylar AtLURE1.2 accumulation. In contrast, the expression of a dominant-negative form of ACTIN8 induced disorganization of the filiform apparatus and loss of polar AtLURE1.2 distribution toward the filiform apparatus. Interestingly, after pollen tube reception, F-actin became unclear for a few hours in the persistent synergid cell, which may be involved in pausing and resuming pollen tube attraction during early polytubey block. Our data propose the central role of F-actin in the maintenance of cell polarity and function of male-female communication in the synergid cell.

## Introduction

Sexual plant reproduction proceeds with multicellular haploid tissues termed gametophytes. Most flowering plants, including *Arabidopsis thaliana,* produce a *Polygonum*-type female gametophyte that contains seven cells: one egg cell, one central cell, two synergid cells and three antipodal cells. On the other hand, pollen grains or pollen tubes are the male gametophytes and contain fewer cells, with a tip-growing pollen tube vegetate cell that contain two sperm cells. Sexual reproduction is accomplished when the egg cell and central cell are fertilized by two sperm cells delivered by the pollen tube. Although the synergid cell itself is not fertilized, it is responsible for almost all communications between female and male gametophytes prior to double fertilization. The synergid cell controls pollen tube attraction, pollen tube reception, termination of pollen tube attraction, and fertilization recovery (Dresselhaus et al., 2016).

In the past two decades, many synergid cell-specific genes have been identified that are required for the communication between the female and male gametophytes. In some species, various pollen tube attractants secreted from the synergid cell have been identified: ZmEA1 in maize (Marton et al., 2005; Marton et al., 2012); LURE1 and LURE2 in *Torenia* (Okuda et al., 2009); AtLURE1s, XIUQIUs, and TICKETs in *A. thaliana* (Takeuchi and Higashiyama, 2012; Zhong et al., 2019; Meng et al., 2019).

Receptor-like kinases on the pollen tube tip, including PRK6, interact with AtLURE1 and control the directional growth of the pollen tube (Takeuchi and Higashiyama, 2016). After arrival of the pollen tube, one of the two synergid cells receives the pollen tube contents, including the two sperm cells. The receptive synergid regulates pollen tube discharge via CrRLK1L-type receptor-like kinases such as FERONIA, ANJEA, and HERCULES RECEPTOR KINASE 1 (HERK1) (Escobar-Restrepo et al., 2007; Galindo-Trigo et al., 2020), a GPI-anchored protein LORELEI (Capron et al., 2008; Tsukamoto et al., 2010), and a Mildew resistance locus o (MLO) family-related protein NORTIA (Kessler et al., 2010). Genetic and biochemical approaches have begun to uncover their cooperative mechanisms during pollen tube reception (Li et al., 2015; Ju et al., 2021). Following pollen tube reception, ovules prevent the attraction of extra pollen tubes, a process called polytubey block, reducing risk of egg cell fertilization by more than one sperm cell known as polyspermy (Kasahara et al., 2012; Beale et al., 2012; Maruyama and Higashiyama, 2016). In early ovular polytubey block, activated just before or upon double fertilization, synergid cells produce nitric oxide (NO) in a FERONIA dependent manner. The NO signal not only inactivates AtLURE1 peptides via post-translational nitrosation, but also inhibits the AtLURE1 secretory system by an unknown mechanism (Duan et al., 2020). In addition, the clearance of attractants may be accelerated by fertilization-dependent secretion of aspartic endopeptidases from the zygote (Yu et al., 2021). On the other hand, later stages of the polytubey block are initiated a few hours after double fertilization by the inactivation of the non-receptive persistent synergid. Representative features of the persistent synergid elimination are nuclear degeneration and cell-to-cell fusion between the persistent synergid and the endosperm (Völz et al., 2013; Maruyama et al., 2013; Maruyama et al., 2015; Pereira et al., 2016). If ovules are not fertilized by the first pollen tube, they pause the degeneration of the persistent synergid and resume the attraction of the next pollen tube (Kasahara et al., 2012; Beale et al., 2012).

The synergid cell displays a highly polarized morphology, characterized by concentration of the protoplasm at the micropylar side, facing toward the pollen tube entry pathway in the ovule, and a large central vacuole occupies at the opposite chalazal side (Mansfield et al., 1991). The micropylar end of the synergid cell has the filiform apparatus, an active communication domain with complex plasma membrane invaginations and thick cell walls, that presumably has pivotal roles in secreting plenty of pollen tube attractant peptides and localizing pollen tube reception factors (Johnson et al., 2019). Synergid cell shape formation and dynamic protein relocation or secretion may be regulated by the cytoskeletons. Immunostaining studies have shown radial microtubules spreading from the filiform apparatus in various species including *A. thaliana* (Webb and Gunning, 1994). In contrast, filamentous actin (F-actin) is widely distributed longitudinally along the micropyle-chalazal axis (Webb and Gunning, 1994). However, the significance of the cytoskeletons in synergid cells remains unclear.

In this study, we aimed to investigate the functions of the cytoskeleton in *A. thaliana* synergid cells. We used genetic and pharmacological approaches to induce microtubule- and actin-depolymerization. Our results indicate that microtubule destruction compromises the elongation of plasma membrane invaginations in the filiform apparatus. On the other hand, disruption of F-actin caused severe disorganization of synergid cell morphology, including phenotypic alterations such as incomplete filiform apparatus formation and aberrant positions of the central vacuole. Furthermore, F-actin destruction impaired the secretion of pollen tube attractant peptides and caused female sterility. Interestingly, the longitudinal F-actin pattern disappeared in the persistent synergid after pollen tube discharge, suggesting dynamic regulation of F-actin during fertilization. Altogether, these findings elucidated flexibility of the F-actin-mediated directional secretion system that maintains morphology and function of the synergid cell.

## RESULTS

### Morphology of filiform apparatus

Filiform apparatus was usually described as a finger-like structure from observations of mature ovules by transmission electron microscopy (Jensen, 1965; Diboll and Larson, 1966; Schulz and Jensen, 1968). To update our knowledge on the filiform apparatus morphology, we conducted three-dimensional (3D) reconstruction of mature *A. thaliana* ovule by focused ion beam-scanning electron microscopy (FIB-SEM) (Oi et al., 2017). A 3D image reconstructed from the serial sections (713 sequential electron micrographs with 25 nm step size) showed a sponge-like porous domain of cell walls at the micropylar end of the filiform apparatus, rather than the finger-like pattern that are often used to depict filiform apparatus in 2D electron micrographs (FA and Po in Figure 1A and 1B; Movie 1). According to the 3D model, the surface area of plasma membrane in the filiform apparatus was 3.7-fold larger than the presumed cell wall area excluding invaginations (72.3 μm^2^ and 19.8 μm^2^, respectively).

**Figure 1.**
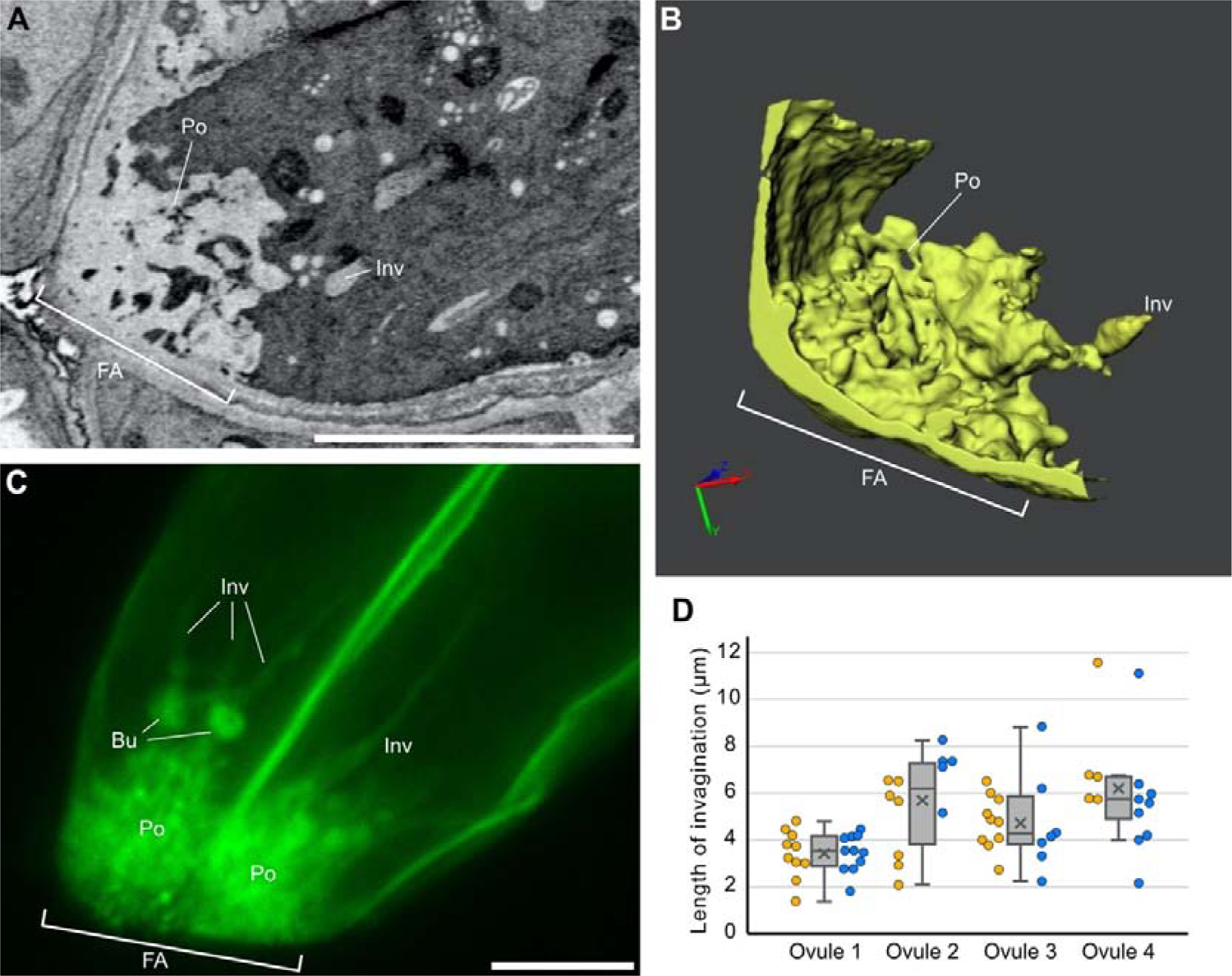
Morphology of filiform apparatus in *Arabidopsis thaliana*. **A, B,** Filiform apparatus analyzed by FIB-SEM (See also Movie 1). A single plane selected from the serial section (A). 3D reconstruction image of the synergid cell wall (B). **C,** Plasma membrane pattern visualized by *pMYB98::3×mNG-SYP132*. A maximum intensity projection from the selected Z-serial sequences in Movie 2 is shown. **D,** Length of plasma membrane invaginations at the filiform apparatus in the *pMYB98::3×mNG-SYP132* plants. Box-and-whisker plots show median (center line), mean (cross mark), upper and lower quartiles (box), maximum and minimum (whiskers), and left and right points corresponding the data from each synergid cell (orange and blue solid circles). Abbreviations: FA, filiform apparatus; Po, porous structure; Inv, invagination; Bu, bulge. Scale bars: 5 μm in (A) and (C).

Next, we generated a plasma membrane marker *pMYB98::3×mNG-SYP132*, expressing Syntaxin of plant 132 (SYP132) tagged with tandem fusions of three mNeonGreen from the synergid cell-specific *MYB98* promoter (Uemura et al., 2004; Shaner et al., 2013; Kasahara et al., 2005) and analyzed complex structures of the filiform apparatus in live synergid cells by confocal microscopy (Figure 1C; Movie 2). Z-series confocal images showed that the porous structure exhibited strong fluorescent signals due to the accumulations of vesicular- or folded-plasma membranes (Figure 1C, Po). In contrast to the FIB-SEM data, we observed long protrusions of the tubular plasma membrane from the porous area toward the inside of the cell (Figure 1C, Inv). Among the eight synergid cells in the four ovules, we observed an average of 8.0±2.3 invaginations per cell with a length ranging from 1.8 to 11.5 μm (Figure 1C). The size of the invagination was comparable between pairs of synergid cells from the same ovule but varied among different ovules (Figure 1D). In addition, bulges were observed along the invaginations in three ovules (Figure 1C, Bu).

### Morphology and dynamics of cytoskeletons in synergid cell

The formation of complex plant cell morphologies often requires F-actin and/or microtubules (Eng and Sampathkumar, 2018). To investigate pattern of these cytoskeletons in the live synergid cell, we generated a double marker line expressing an F-actin binding Lifeact peptide tagged with mRUBY3 (Lifeact-mRUBY3) and TUBULIN ALPHA 5 fused with Citrine (Citrine-TUA5) from the synergid cell-specific *MYB98* promoter (Riedl et al., 2008; Shaner et al., 2013; Kasahara et al., 2005). In the mature ovules, mRUBY3 signal of F-actin extended from the micropylar end, where the filiform apparatus resides, and reached beyond synergid nuclei and large central vacuoles to the chalazal end (Figure 2A, left panel). Citrine-labeled microtubules were also longitudinally aligned around the filiform apparatus, forming a radially extended cat-whisker pattern (Figure 2A, middle panel), where F-actin often associated or co-localized with microtubules (Figure 2A, right panel).

**Figure 2.**
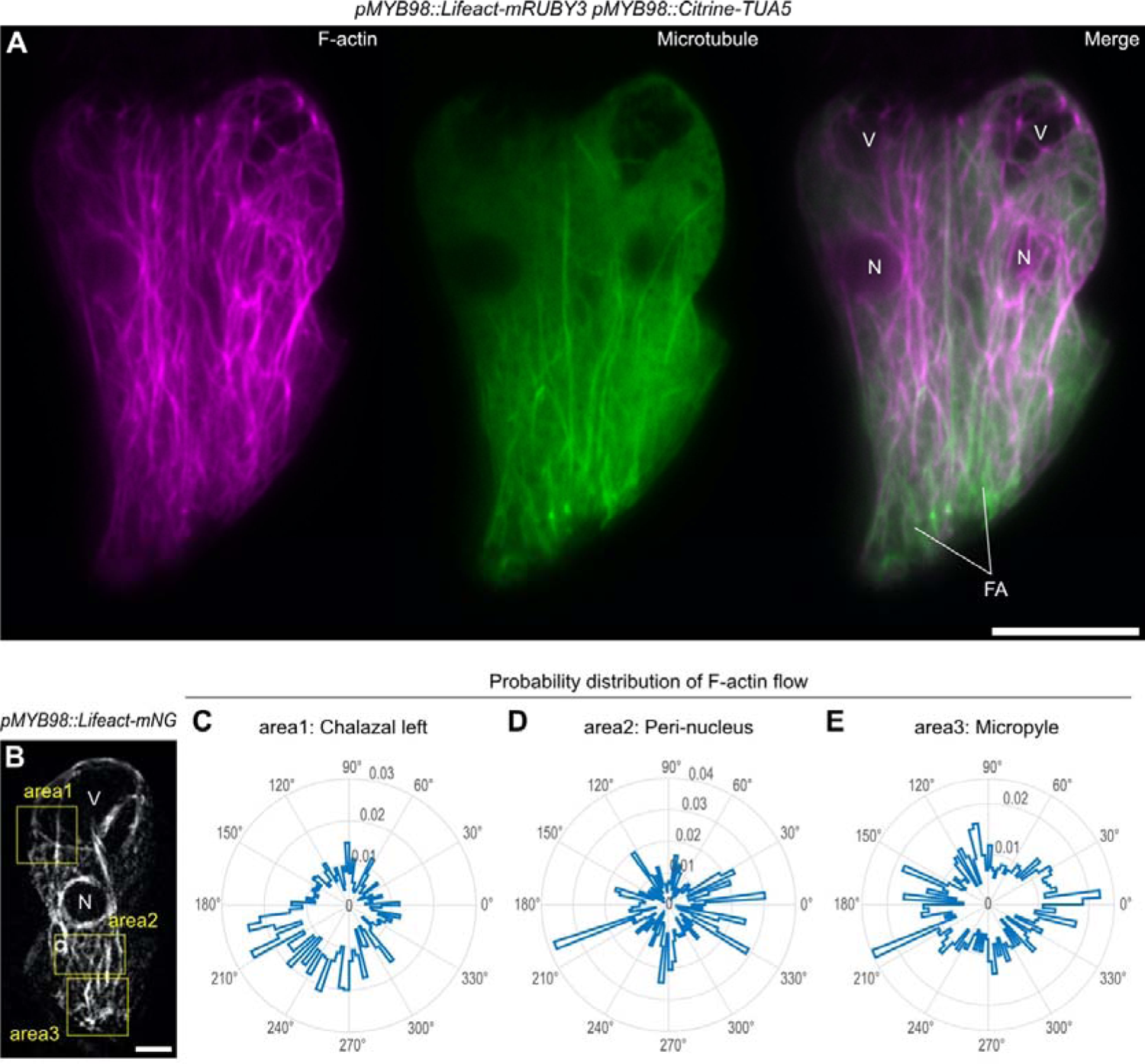
Distributions of cytoskeletons in synergid cells. **A,** Maximum intensity projection confocal images of a pair of synergid cells expressing *Lifeact-mRUBY3* F-actin marker (Magenta) and *Citrine-TUA5* microtubule marker (Green) from the *MYB98* promoter. **B,** The first frame of time-lapse imaging of F-actin dynamics in the synergid cell. A mature ovule in the transgenic plant expressing *Lifeact-mNeonGreen* F-actin marker (Grey) from the *MYB98* promoter (*pMYB98::Lifeact-mNG*) was captured in 30 sec intervals. The yellow boxes indicate the areas where F-actin flow was analyzed as shown in (C-E). V, vacuole; N, nucleus; FA, filiform apparatus; Scale bars, 10 μm in (A) and (B). **C-E,** Radar plots showing probability distributions of the directions of F-actin flow vectors in each area (the yellow box in (B)) obtained via spatiotemporal image correlation spectroscopy analysis. The probability was calculated from cumulative frequency of the F-actin flow vector angles obtained from the entire frames of time-lapse imaging. The probability values are scaled in the radial axes on the radar plots.

In the egg cell and central cell, F-actin is generated at the cell periphery and migrates toward the nucleus, supporting the rapid merger of female and male nuclei (karyogamy) after gamete fusions (plasmogamy) (Kawashima et al., 2014; Ohnishi et al., 2014; Ohnishi and Okamoto, 2015). To examine whether fertilization-incompetent gametophytic cells (*i.e.,* synergid cells) have the similar nucleopetal F-actin dynamics, we observed F-actin dynamics in the synergid cells of mature ovules (*pMYB98::Lifeact-mNG*) by time-lapse imaging at 30 sec intervals. The movie showed that actin cables actively moved longitudinally (Movie 3), and we further performed the spatiotemporal image correlation spectroscopy analysis (Ashdown et al., 2015) to evaluate directions of F-actin flow at four areas (two in the chalazal and one each in peri-nuclear and micropylar regions). Radar plots indicated that the directions of F-actin flow were mainly from chalazal to micropyle, except for the micropylar region (Figure 2B-2E and S2). Around the filiform apparatus, the F-actin flow was omnidirectional, implying active rearrangements of F-actin. These results show that, unlike the nucleopetal F-actin dynamics of the gametes, overall F-actin dynamics in the synergid cell are toward the filiform apparatus.

### Genetic disruption of microtubule formation

To analyze the roles of microtubules in the synergid cell, we used PHS1ΔP, a dominant mutant of PROPYZAMIDE-HYPERSENSITIVE 1 (PHS1). PHS1 is an α-tubulin kinase consisting of a central tubulin kinase domain and a C-terminal phosphatase domain (Fujita et al., 2013). PHS1ΔP lacks the C-terminal phosphatase domain and shows high tubulin kinase activity, inducing microtubule depolymerization via overproduction of polymerization-inefficient phosphorylated α-tubulins (Fujita et al., 2013). We generated two transgenic lines homozygous for a transgene expressing the *PHS1ΔP* from the *MYB98* promoter (*pMYB98::PHS1ΔP*) that were also homozygous for two reporter genes visualizing synergid cell morphology: the *pRPS5A::H2B-tdTomato* (*RHT*) and *pES2::3×mNG-SYP132* (ENS). *RHT* is a nuclear marker expressing tdTomato-tagged HISTONE H2B from the ubiquitously active *Ribosomal Protein Subunit 5A* promoter (Maruyama et al., 2013). As shown in Figure S1, *ENS* is a plasma membrane marker expressing 3×mNeonGreen-SYP132 from the embryo sac-specific *ES2* promoter (Yu et al., 2005; Pagnussat et al, 2007).

The mature ovules from the *pMYB98::PHS1ΔP;RHT;ENS* transgenic lines were compared with those from the wild-type *RHT;ENS* line using confocal microscopy. In wild-type *RHT;ENS*, 97.7% of the ovules contained the normal seven-celled female gametophyte with two synergid cells, an egg cell, a central cell, and three antipodal cells (Figure 3A, n = 386). On the other hand, in *pMYB98::PHS1ΔP;RHT;ENS*, only 9.3– 12.9% of the gametophytes showed normal cell structure (Figure 3B, Two synergids; Figure 3F), and the remaining ovules exhibited varieties of aberrance. Specifically, 37.2–37.5% of the ovules contained a single synergid cell with either one or two nuclei (Figure 3C, single synergid). Because the *MYB98* promoter becomes active prior to cellularization of the eight-nucleate female gametophyte (Kasahara et al., 2005; Susaki et al., 2021), these single synergid ovules were likely generated due to the inadequate spindle and/or cell plate formation caused by PHS1ΔP-mediated microtubule depolymerization (Figure 3C and 3F, SY-SY fusion). Consistently, we observed more severe phenotypes in *pMYB98::PHS1ΔP;RHT;ENS*; including the disappearance of mNeonGreen-labeled plasma membrane accumulation, a hallmark of the filifor apparatus, in 49.5–53.6% of the ovules (Figure 3D and 3E, No FA). Occasionally, these ovules contained no nuclei in the putative synergid region while the central cell had multiple nuclei (Figure 3E, 9.5–10.3%). These phenotypes suggest that the absence of the filiform apparatus may reflect the failure of synergid cell differentiation due to incomplete cytokinesis between the two synergid cells and the central cell (Figure 3D– 3F, CC-SY fusion).

**Figure 3.**
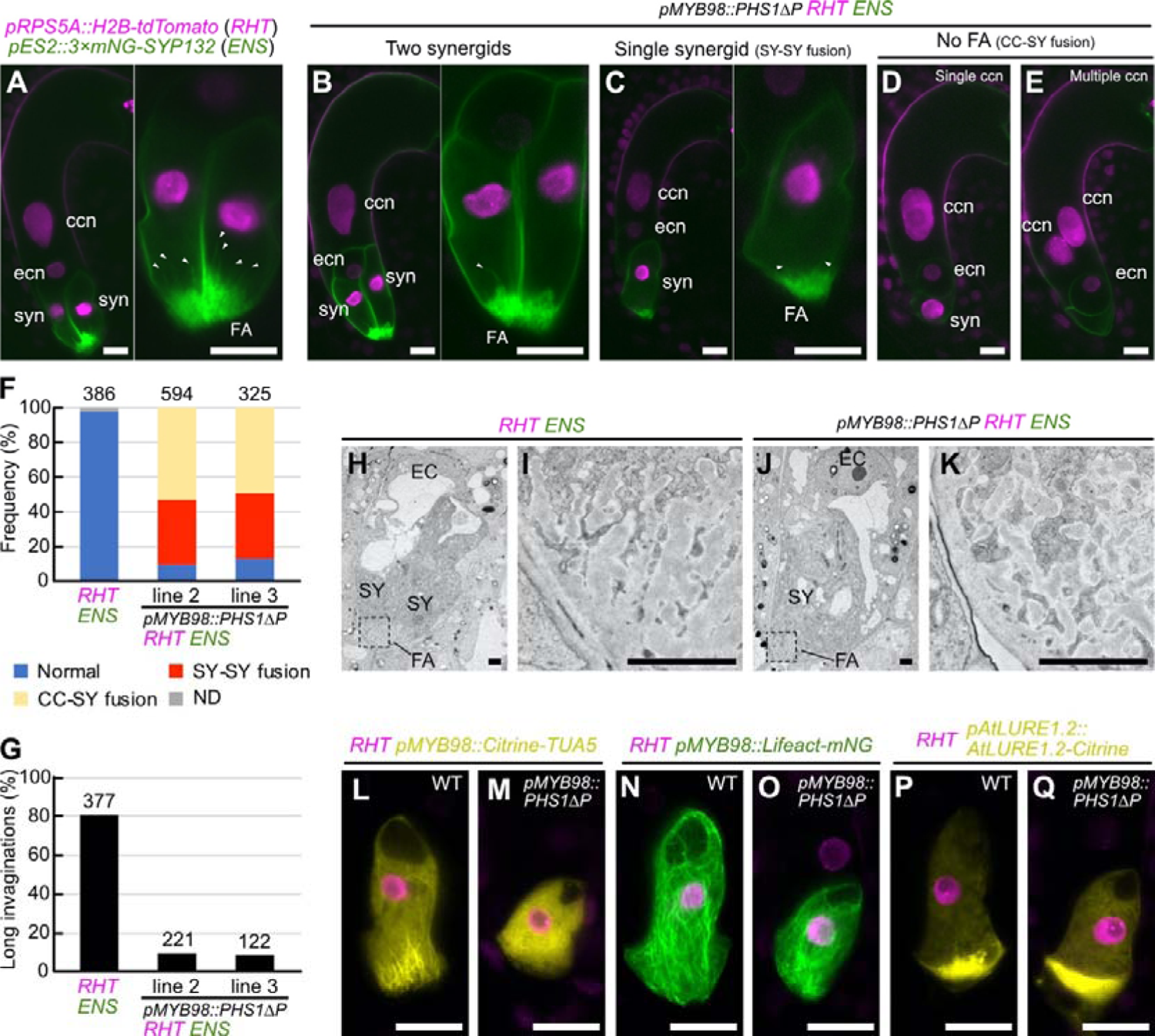
Microtubule is required for the growth of plasma membrane invaginations at the filiform apparatus. **A,** Wild-type ovule carrying the nuclear marker (*RHT*) and plasma membrane marker (*ENS*). **B–E,** Ovules containing two synergid cells (B), single synergid cell (C), a single central cell nucleus without filiform apparatus (D), or multiple central cell nuclei without filiform apparatus (E) observed in the *RHT;ENS* double marker line that were also homozygous for the microtubule destruction gene *PHS1ΔP* expressed from the *MYB98* promoter (*pMYB98::PHS1ΔP*). In (A–C), the synergid cell(s) in the left panel is magnified and shown in the right panel. **F, G,** Frequency of normal and aberrant female gametophyte (F) and percentages of filiform apparatus with long invaginations (G) in wild-type *RHT;ENS* line and two *pMYB98::PHS1ΔP;RHT;ENS* lines. **H–K,** Transmission electron micrographs of wild-type ovule in the *RHT;ENS* line (H) and single synergid cell-containing ovule of the *pMYB98::PHS1ΔP;RHT;ENS* line (J). Magnification of areas of the filiform apparatus indicated with dashed boxes in (H) and (J) were shown in (I) and (K), respectively. **L–Q,** Confocal images of synergid cells expressing *RHT* nuclear marker and other reporter genes showing synergid cell structures in wild-type plants (L, N, P) and *pMYB98::PHS1ΔP* transgenic lines (M, O, Q). The synergid cell-specific reporters were as follows: *pMYB98::Citrine-TUA5* (microtubules; L, M), *pMYB98::Lifeact-mNG* (F-actin; N, O), *pAtLURE1.2::AtLURE1.2-Citrine* (pollen tube attractant; P, Q). Abbreviations: *RHT*, *pRPS5A::H2B-tdTomato*; *ENS*, *pES2::3×mNG-SYP132*; ccn, central cell nucleus; ecn, egg cell nucleus; syn, synergid cell nucleus; FA, filiform apparatus; SY, synergid cell; EC, egg cell; CC, central cell. Scale bars: 20 μm in (A) to (E) and (L) to (Q); 2 μm in (H) to (K).

In the following experiments, we focused on the single-synergid ovules because they were observed more frequently than the two-synergid ovules (Figure 3C and 3F). Around the filiform apparatus, only ∼10.0% of the single-synergid ovules produced long plasma membrane invaginations (Figure 3C and 3G, greater than ∼6 µm), which was considerably fewer than that of normal synergid cells in the wild-type *RHT;ENS* ovules (Figure 3A and 3G, 80.9%, n = 377). Next, we performed transmission electron microscopy to analyze the cell walls at the filiform apparatus (Figure 3H-3K). In wild-type *RHT;ENS*, the cell walls exhibited typical finger-like pattern consisting of an electron dense central region surrounded by a translucent peripheral layer (Figure 3H and 3I, 100%, n = 7). Similar finger-like cell walls were also observed in the single-synergid ovules of *pMYB98::PHS1ΔP RHT;ENS* (Figure 3J and 3K, 80%, n = 5).

To investigate the expression and localization of the cytoskeletons and pollen tube attractant, we introduced the microtubule, F-actin, and pollen tube attractant markers into the wild-type *RHT* and *pMYB98::PHS1ΔP;RHT* lines (Figure 3L-3Q). The single-synergid ovules carrying *pMYB98::PHS1ΔP* displayed no fibers, visualized using the *pMYB98::Citrine-TUA5* microtubule marker, confirming the significant effect of PHS1ΔP expression microtubule depolymerization (Figure 3L and 3M). Normal longitudinal F-actin *(pMYB98::Lifeact-mNeonGreen)* was observed in the single-synergid ovules of the *pMYB98::PHS1ΔP* line (Figure 3N and 3O). Finally, the pollen tube attractant marker, *pAtLURE1.2::AtLURE1.2-Citrine*, showed a comparable Citrine focus at the filiform apparatus in both wild-type and *pMYB98::PHS1ΔP* lines, indicating of normal polarized secretion in these ovules (Figure 3P and 3Q). Altogether, these results suggest that whisker-like microtubules facilitate the growth of plasma membrane invaginations but have a limited effect on the formation of the basal structure of the filiform apparatus and cell polarity.

To investigate pollen tube attraction and fertility, pistils from the wild-type *RHT;ENS* and *pMYB98::PHS1ΔP;RHT;ENS* lines were pollinated with the non-transgenic Col-0 pollen (Figure 4). At one-day-after pollination (1 DAP), pollen tubes were visualized by CongoRed staining. In the wild-type *RHT;ENS* pistils, 97.5% of the ovules received pollen tubes and displayed nuclear proliferation in the endosperm, showing successful pollen tube attraction and fertilization (Figure 4A, n = 360). Similar fertilization phenotypes were also observed in 49.7% of the ovules from *pMYB98::PHS1ΔP;RHT;ENS* plants (Figure 4B, n = 338), which corresponded to the total frequency of the single-synergid and two-synergid ovules (Figure 3B, 3C, 3F). The remaining 50.3% did not proliferate the endosperm nuclei (Figure 4C-4E and 4G), implying fertilization abnormalities in these ovules, which also showed no plasma membrane accumulation at the filiform apparatus (Figure 3D–F). In the *pMYB98::PHS1ΔP;RHT;ENS* plants, normal seed development was observed in ∼70% at 10 DAP (Figure 4H-4J), which were comparable to the ovules receiving a pollen tube at 1 DAP (Figure 4F). Thus, the subset of the PHS1ΔP-expressing ovules might show fertilization delay soon after pollen tube reception (Figure 4C and 4F, 22,2%) and would be gradually fertilized by 8 DAP. Taken together, PHS1ΔP-mediated microtubule depolymerization and the loss of long invaginations at the filiform apparatus would not abolish pollen tube attraction and reception by the synergid cell.

**Figure 4.**
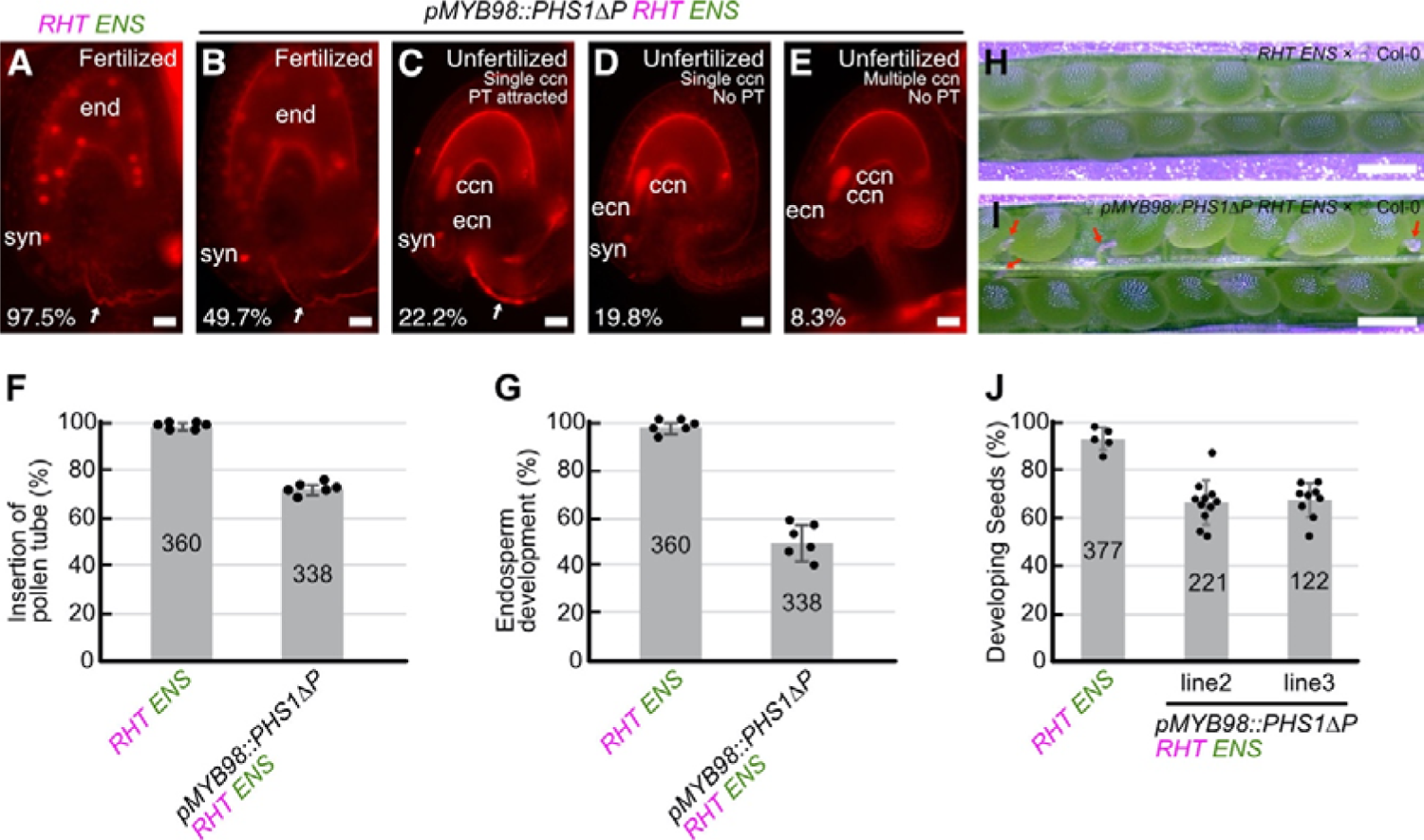
Fertilization of ovules expressing PHS1ΔP in synergid cells. **A–E,** Analysis of pollen tube attraction and seed development in wild-type plant carrying the nuclear marker (*RHT*) and plasma membrane marker (*ENS*) (A), and *pMYB98::PHS1ΔP;RHT;ENS* transgenic plant (B–E). Ovules were dissected from pistils at one day after pollination with wild-type Col-0 pollen and observed after CongoRed staining. Frequency of each ovule phenotype was shown in the bottom-left. White arrows indicate pollen tubes. **F, G,** Percentages of ovules displaying pollen tube insertions (F) and endosperm development (G). **H, I,** Representative images of developing seeds in pistils from wild-type *RHT;ENS* plant (H), and *pMYB98::PHS1ΔP;RHT;ENS* transgenic line (I), analyzed at eight days after pollination with wild-type Col-0 pollen. Red arrows indicate undeveloped ovules. **J,** Frequencies of developing seeds analyzed in (H) and (J). Abbreviations: *RHT*, *pRPS5A::H2B-tdTomato*; *ENS*, *pES2::3×mNG-SYP132*; ccn, central cell nucleus; ec n, egg cell nucleus; syn, synergid cell nucleus; end, endosperm nuclei. Scale bars: 20 μm in (A) to (E); 0.5 mm in (H) and (J).

### Expression of a dominant-negative actin in the synergid cell caused severe defect in pollen tube attraction

To examine the roles of F-actin in synergid cell, we generated transgenic plants expressing wild-type ACTIN8 (*WT-ACTIN*) or the dominant-negative form of ACTIN8 (*DN-ACTIN*) from the *MYB98* promoter. Homozygous plants were not recovered in four independent *pMYB98::DN-ACTIN* lines, implying reduced fertility in the female gametophyte. To further investigate that, we compared 8 DAP seeds in pistils from Col-0 wild-type plants and *pMYB98::DN-ACTIN* hemizygous mutants pollinated with Col-0 wild-type pollen (Figure 5A-5C). Wild-type pistils contained almost full sets of seeds (Figure 5A and 5C). However, approximately half of the ovules remained undeveloped in the *pMYB98::DN-ACTIN* line (Figure 5B and 5C). Next, we analyzed the hygromycin resistance in these F_1_ siblings. Hygromycin resistance in the *pMYB98::WT-ACTIN* hemizygous line was 52.7%, suggesting normal inheritance in the *pMYB98::WT-ACTIN* plants from the female gametophyte (Figure 5D). Nevertheless, hygromycin resistance in the four *pMYB98::DN-ACTIN* hemizygous lines ranged from 2.8 to 9.5% (Figure 5D). When wild-type Col-0 pistils were pollinated with pollen from the *pMYB98::WT-ACTIN* or *pMYB98::DN-ACTIN* hemizygous plants, no reproductive abnormalities, such as reduced seed set or decreased hygromycin resistance, were observed in the F_1_ siblings (Figure 5C and 5E). Taken together, we concluded that the *pMYB98::DN-ACTIN* caused sterility exclusively in the female gametophyte.

**Figure 5.**
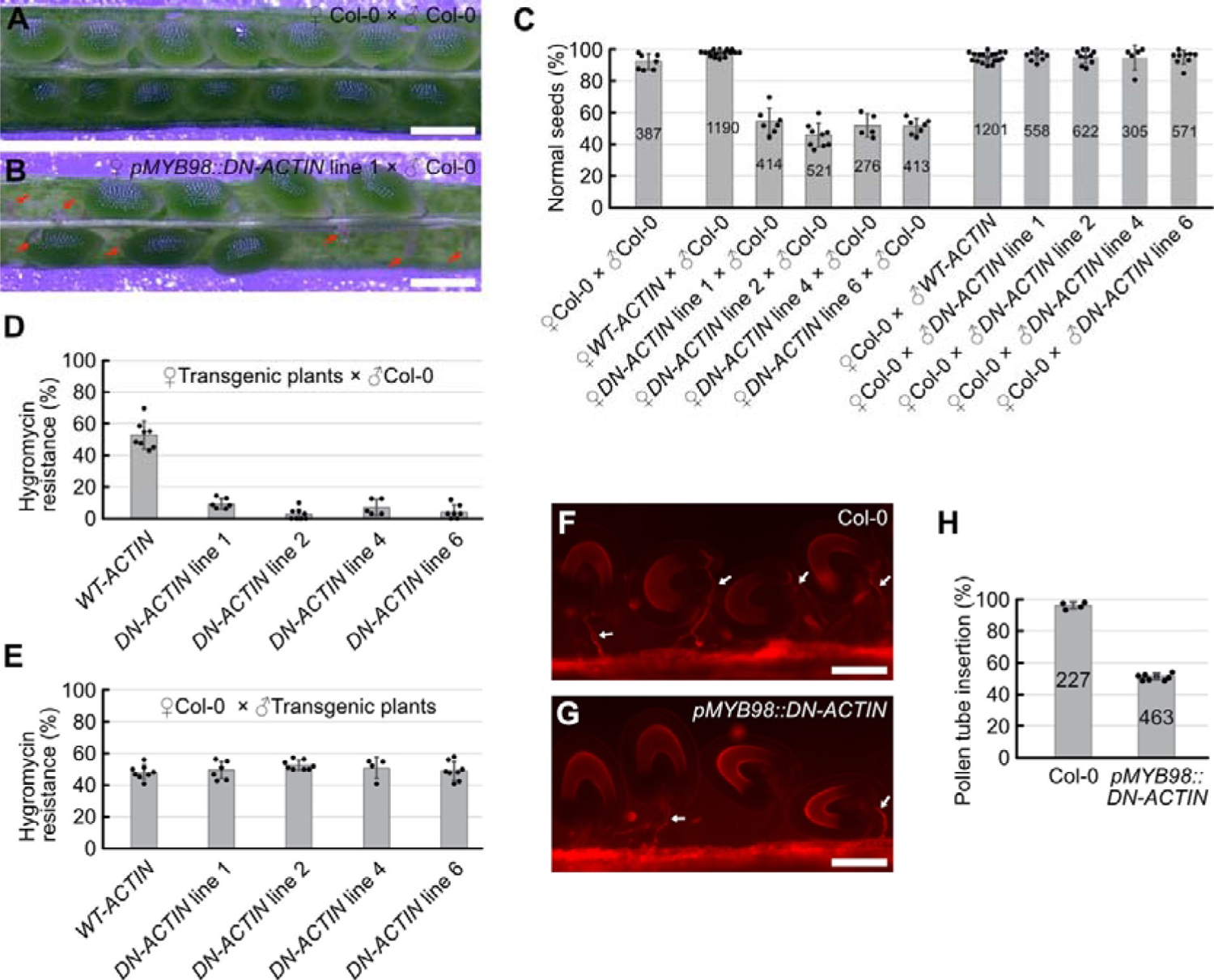
F-actin is required for pollen tube attraction. **A, B,** Representative images of developing seeds in pistils from wild-type Col-0 plants (A), and *pMYB98::DN-ACTIN* hemizygous lines (B), analyzed at eight days after pollination with wild-type Col-0 pollen. Red arrows indicate undeveloped ovules. **C,** Percentages of normal seeds harvested from pistils after various combinations of cross-pollination among wild-type Col-0 plants, *pMYB98::WT-ACTIN* hemizygous plants (*WT-ACTIN*), and *pMYB98::DN-ACTIN* hemizygous plants (*DN-ACTIN*). **D, E,** Transmissions of *pMYB98::WT-ACTIN* and *pMYB98::DN-ACTIN* transgenes via female gametophyte (D) or male gametophyte (E). Gene transmission was analyzed by frequencies of hygromycin-resistant F1 siblings generated by cross-pollinations in (B). **F, G,** Pollen tube growth pattern in pistils from wild-type Col-0 (F) and *pMYB98::DN-ACTIN* hemizygous (line 1) (G) plants. Pistils were pollinated with wild-type Col-0 pollen and the pollen tubes were visualized by a CongoRed staining at one day after pollination. White arrows indicate pollen tubes. **H,** Percentages of pollen tube-inserted ovules analyzed in (F) and (G). Scale bars: 0.5 mm in (A) and (B); 100 μm in (F) and (G).

To further investigate the cause of female sterility, we first examined pollen tube attraction. Pistils from wild-type and *pMYB98::DN-ACTIN* plants were pollinated with wild-type Col-0 pollen, and the pollen tube growth pattern was analyzed at 1 DAP using CongoRed staining. In wild-type pistils, 96.2% of the ovules received the pollen tube, indicative of successful pollen tube attraction (Figure 5F and 5H, n = 227), whereas pollen tube attraction was observed only in 51.3% of the ovules from the *pMYB98::DN-ACTIN* line (Figure 5G and 5H, n = 463). Therefore, *pMYB98::DN-ACTIN* may reduce ovule fertility through a pollen tube guidance defect.

### Loss of F-actin in the synergid cell impairs secretion of pollen tube attractant peptides

The pollen tube attraction defect observed in *pMYB98::DN-ACTIN* may be explained by a decrease in the secretion of pollen tube attractant peptides from the synergid cell. To assess this hypothesis, we compared the immunostaining pattern of AtLURE1.2 in mature ovules from non-transgenic Col-0, *myb98* homozygous, and *pMYB98::DN-ACTIN* hemizygous plants (Figure S3). AtLURE1.2 signals were detected on the surface of the micropyle and/or funiculus in ∼90% of the ovules from non-transgenic Col-0 plants (Figure S3A; n = 206). The AtLURE1.2 signal was below the detection level in the *myb98* homozygous mutant, as reported previously (Figure S3B; n = 231; Takeuchi and Higashiyama, 2012). In the *pMYB98::DN-ACTIN* hemizygous plant, AtLURE1.2 signals were only observed in ∼40% of the ovules (Figure S3C; n = 260), indicating a reduction of AtLURE1.2 secretion in the *DN-ACTIN* expressing ovules. To further examine pollen tube attractants in the synergid cells, we used a translational fusion reporter of *AtLURE1.2* and *Citrine* (*pAtLURE1.2::AtLURE1.2-Citrine*). Consistent with the previous report, the fluorescent signal of AtLURE1.2-Citrine was specifically observed in the synergid cell with the strongest signal detected at the filiform apparatus (97.0%, n = 339; Figure 6A, 6B and 6G; Takeuchi and Higashiyama, 2012). In the *pMYB98::DN-ACTIN* hemizygous plants that were homozygous for the *pAtLURE1.2::AtLURE1.2-Citrine*, Citrine signal was detected in 92.9% of ovules (n = 432; Figure 6C-6D), suggesting that DN-ACTIN had little or no effect on cell differentiation and gene expression in the synergid cell. However, a normal AtLURE1.2-Citrine secretion pattern was observed only in 46.3% of ovules (WT; Figure 6C, 6D and 6G). In the remaining ovules that were expected to carry *pMYB98::DN-ACTIN*, AtLURE1.2-Citrine did not accumulate at the filiform apparatus but rather was uniformly distributed in the cytoplasm (*pMYB98::DN-ACTIN*; Figure 6E and 6F). Abnormalities were also observed in the central vacuole. Wild-type ovules presented single large central vacuole at the chalazal side (Figure 6B and 6D), whereas ovules from *pMYB98::DN-ACTIN* hemizygous line often contained multiple small vacuoles located in the middle or micropylar side of the synergid cell (Figure 6F). In agreement, similar results were obtained with the pollen tube attractant marker, *XIUQIU4-Citrine* (Figure S4). These data indicate the aberrant secretion of AtLURE1.2 and XIUQIU4 in the DN-ACTIN expressing synergid cells.

**Figure 6.**
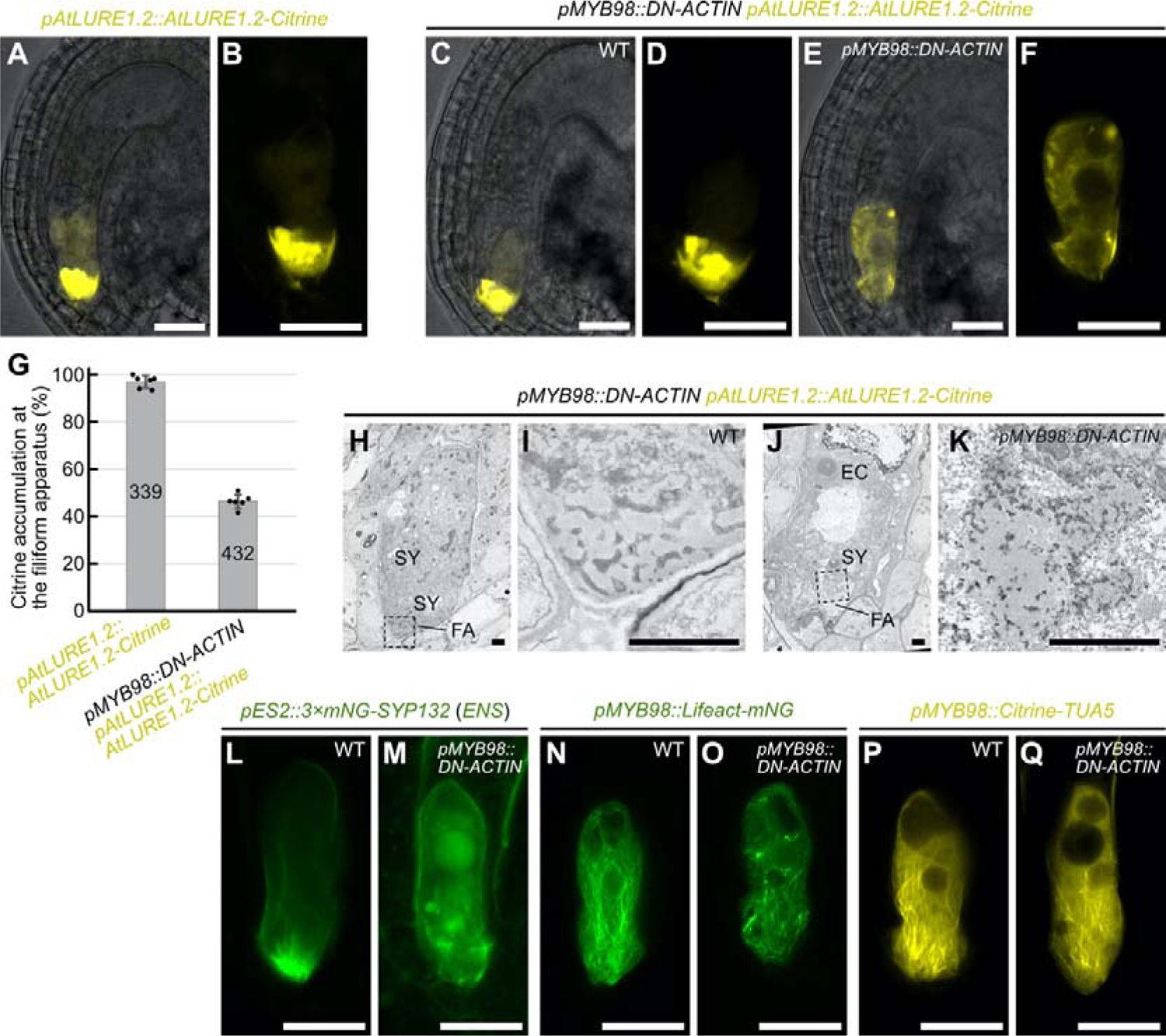
F-actin is necessary for directional protein secretion in synergid cell. **A–F,** Fluorescent signals of *pAtLURE1.2::AtLURE1.2-Citrine* pollen tube attractant marker in wild-type (A, B), and *pMYB98::DN-ACTIN* hemizygous (line 1) (C–F) plants. Merges of fluorescent images and differential interference contrast images were shown in (A), (C), and (E). Fluorescent images of magnified synergid cells were shown in (B), (D), and (F). The *pMYB98::DN-ACTIN* hemizygous plant showed segregation of wild-type ovules with normal fluorescent pattern (C, D), and abnormal ovules with little AtLURE1.2-Citrine signal at the filiform apparatus (E, F). **G,** Frequency of ovules displaying normal AtLURE1.2-Citrine accumulation at the filiform apparatus analyzed in (A) to (F). **H–K,** Transmission electron micrographs of wild-type (H, I) and abnormal (J, K) ovules segregated in pistils from a *pAtLURE1.2::AtLURE1.2-Citrine* transgenic plant hemizygous for the *pMYB98::DN-ACTIN* (line 1). Magnification of filiform apparatus indicated by dashed boxes in (H) and (J) were shown in (I) and (K), respectively. **L–Q,** Confocal images of synergid cells expressing plasma membrane marker (*pES2::3×mNG-SYP132*; L, M), F-actin marker (*pMYB98::Lifeact-mNG*; N, O), and microtubule marker (*pMYB98::Citrine-TUA5*; P, Q) in the *pMYB98::DN-ACTIN* hemizygous plants (line 1). Representative images of wild-type ovules (L, N, P) and aberrant ovules (M, O, Q) were shown. Abbreviations: FA, filiform apparatus; SY, synergid cell; EC, egg cell. Scale bars: 20 μm in (A) to (F) and (L) to (Q); 2 μm in (H) to (K).

We next analyzed the ultrastructure of normal and abnormal ovules segregated from *pMYB98::DN-ACTIN* hemizygous plants that were homozygous for *pAtLURE1.2::AtLURE1.2-Citrine* (Figure 6H-6K). Although normal ovules with a strong Citrine signal at the filiform apparatus showed finger-like cell walls (Figure 6H and 6I; n = 7), abnormal ovules accumulated electron-dense cell wall materials with chaotic pattern (Figure 6J and 6K; n = 5). To examine other morphological aspects of the abnormal synergid cell, marker genes for the plasma membrane and cytoskeletons were introduced into *DN-ACTIN* expressing lines. In the *pMYB98::DN-ACTIN* hemizygous plants that were homozygous for *pES2::3×mNG-SYP132* (*ENS*), 48.6% of the ovules exhibited a normal filiform apparatus pattern, characterized by the accumulation and invagination of mNeonGreen-labeled plasma membrane at the filiform apparatus (Figure 6L; n = 436). The remaining ovules (51.4%) did not have mNeonGreen focus at the micropylar end, and vesicles of various sizes were detected inside of the synergid cell (Figure 6M), suggesting disorganization of the filiform apparatus in the *pMYB98::DN-ACTIN* line. In the *pMYB98::DN-ACTIN* hemizygous plants that were homozygous for *pMYB98::Lifeact-mNeonGreen*, 49.3% of the ovules produced longitudinally aligned F-actin as wild-type ovules observed in Figure 2 (Figure 6M; n = 319), while the remaining ovules showed fragmented F-actin and destruction of the actin cable alignment (Figure 6M). On the other hand, typical whisker-like microtubule extensions at the filiform apparatus did not change between normal and abnormal ovules in the *pMYB98::DN-ACTIN* hemizygous plants that were homozygous for the *pMYB98::Citrine-TUA5* (Figure 6P and 6Q; n = 212). The Citrine-TUA5 pattern indicates independence of F-actin formation and microtubule formation in the synergid cell.

The analyses of the *pMYB98::DN-ACTIN* transgenic lines revealed important roles of F-actin in pollen tube attraction. However, this genetic approach could not clarify whether the pollen tube attraction defect was induced by the loss of the filiform apparatus or inhibition of attractant peptide secretion. Thus, we examined changes in synergid cell morphology within a few hours after actin polymerization inhibition by Latrunculin A (LatA) treatment. First, we confirmed that half-an-hour treatment with 100 μM LatA diminished F-actin in synergid cell from the *pMYB98::Lifeact-mNG* line (Figure 7A and 7B). Interestingly, *pMYB98::Citrine-TUA5* synergid cell displayed normal whisker-like microtubule extensions after 100 μM LatA treatment (Figure 7C and 7D), indicating independent regulation of F-actin and microtubules in the synergid cell. In addition, plasma membrane accumulation and invaginations at the filiform apparatus visualized with *pES2::3×mNG-SYP132* were not altered even 4h after 100 μM LatA treatment (Figure 7E and 7F). Nevertheless, drastic alterations were observed in the *pAtLURE1.2::AtLURE1.2-Citrine* line; which showed a Citrine signal focused at the filiform apparatus in control DMSO-treated ovules that disappeared 4h after 100 μM LatA treatment, which also induced the appearance of smaller vacuoles (Figure 7G and 7H). The LatA-mediated AtLURE1.2-Citrine signal alterations resembled the phenotypes induced by the genetic F-actin destruction in the *pMYB98::DN-ACTIN* lines (Figure 6E and 6F), suggesting that F-actin destruction quickly stalls AtLURE1.2-Citrine secretion in synergid cell even in the presence of a functional filiform apparatus. A similar analysis was performed by treating *pAtLURE1.2::AtLURE1.2-Citrine*, *pES2::3×mNG-SYP132*, *pMYB98::Lifeact-mNG*, and *pMYB98::Citrine-TUA5* lines with the microtubule polymerization inhibitor, Oryzalin (Figure S5). However, an altered fluorescent pattern was observed only with the *pMYB98::Citrine-TUA5* microtubule marker (Figure S5C and S5D). Taken together, these results suggest that F-actin regulates the polarized secretion of pollen tube attractant peptides at the filiform apparatus.

**Figure 7.**
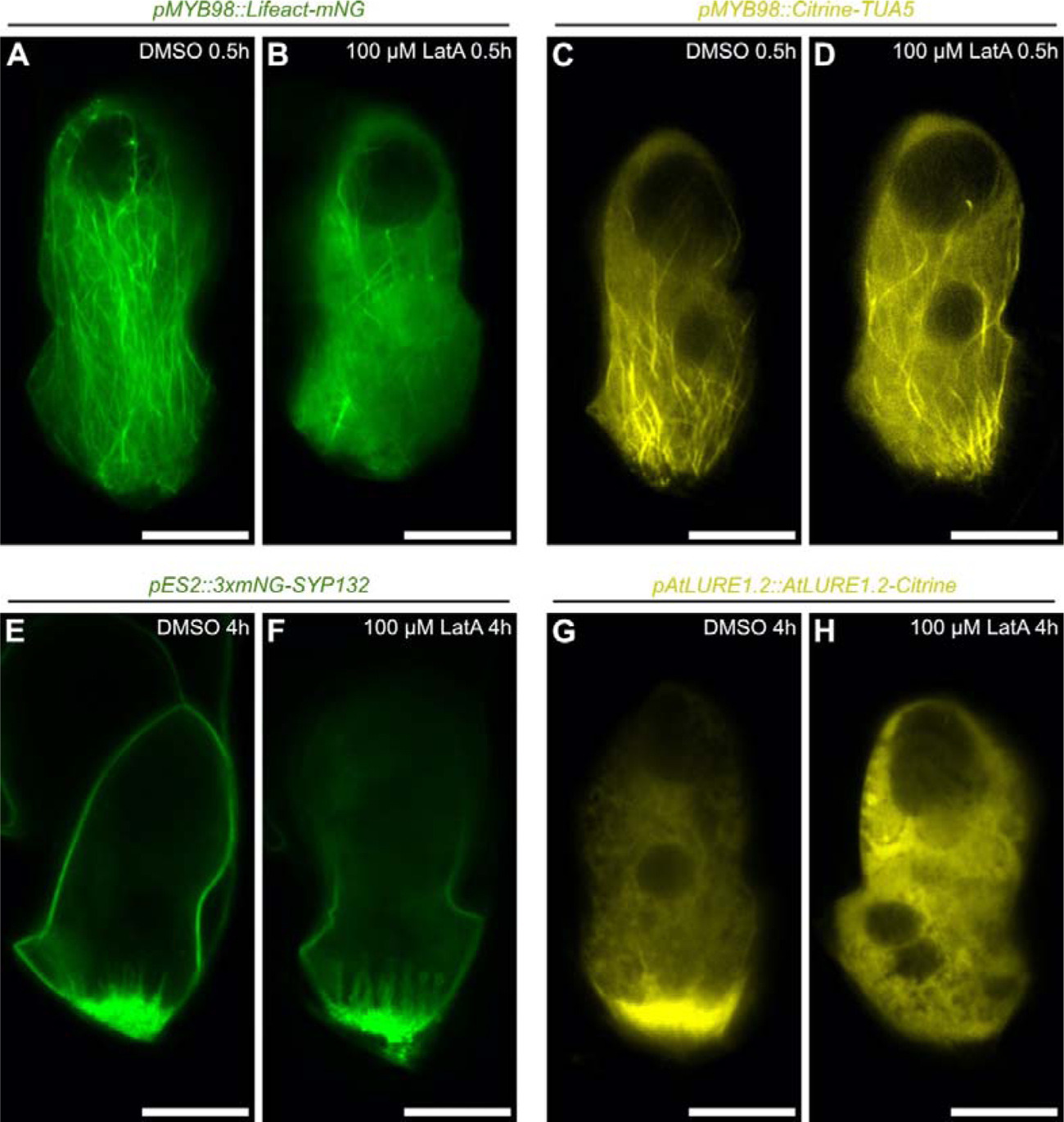
Transient actin depolymerization inhibits AtLURE1.2 accumulation without degeneration of the filiform apparatus. **A–H,** Confocal images of synergid cells in the *pMYB98::Lifeact-mNG* (F-actin marker line) (A, B), *pMYB98::Citrine-TUA5* (microtubule marker line) (C, D), *pES2::3×mNG-SYP132* (plasma membrane marker line) (E, F), and *pAtLURE1.2::AtLURE1.2-Citrine* (pollen tube attractant marker line) (G, H), cultured in a control medium containing 0.4% DMSO (A, C, E, G), or a medium containing 0.4% DMSO and 100 μM Latrunculin A (LatA) (B, D, F, H). Duration of the ovule culture is shown in the top-right. Scale bars: 20 μm.

### Temporal disappearance of longitudinal F-actin in persistent synergid cell after pollen tube discharge

Cumulative evidence supporting F-actin-mediated pollen tube attraction prompted us to investigate F-actin pattern in the persistent synergid during fertilization because it may be involved in the cessation of pollen tube attraction observed after successful double fertilization (Kasahara et al., 2012; Beale et al., 2012). Pistils from *pMYB98::Lifeact-mNG* were pollinated with pollens from a sperm cell-specific nuclear marker *pHTR10::HTR10-mRFP1* (Ingouff et al., 2007), and ovules were analyzed by spinning disk confocal microscopy at 6 h-after-pollination (6 HAP) (Figure 8A-8I). In the unfertilized ovules, normal longitudinal F-actin pattern was observed in both intact synergid cells on different focal planes (Figure 8A, 8E, and 8I). By contrast, different F-actin pattern was observed in two synergid cells after pollen tube reception. According to the sperm nuclei-derived mRFP1 signals, fertilization states of these ovules were classified into three categories: before plasmogamy, before karyogamy, and after karyogamy. Ovules before plasmogamy contained sperm cells closely apposed each other between the egg cell and central cell (Figure 8B and 8F). In the ovules before karyogamy, each separated sperm nucleus was observed in the egg cell and central cell (Figure 8C and 8G). In ovules after karyogamy, the sperm-derived mRFP1 signal was dispersed in the zygote nucleus or primary endosperm nucleus by chromatin decondensation in ovules after karyogamy (Figure 8D and 8H). Throughout these fertilization stages, actin cables were unclear in the receptive synergid and the fluorescent proteins leaked into and marked the boundary of the egg cell and central cell (Figure 8B-8D). In addition, we often observed mNeonGreen foci around the chalazal end of the receptive synergid, which may correspond to the “actin corona” visualized by Rhodamine-phalloidin staining in *Nicotiana tabacum* ovules (Russell, 1993; Huang and Russell, 1994). When the pistils from *pMYB98::Lifeact-mNG* were fertilized by the tagRFP-expressing pollen tubes (*pLAT52::tagRFP*), we found moderate colocalization of cytosolic tagRFP with the actin corona-like structure (Figure S6).

**Figure 8.**
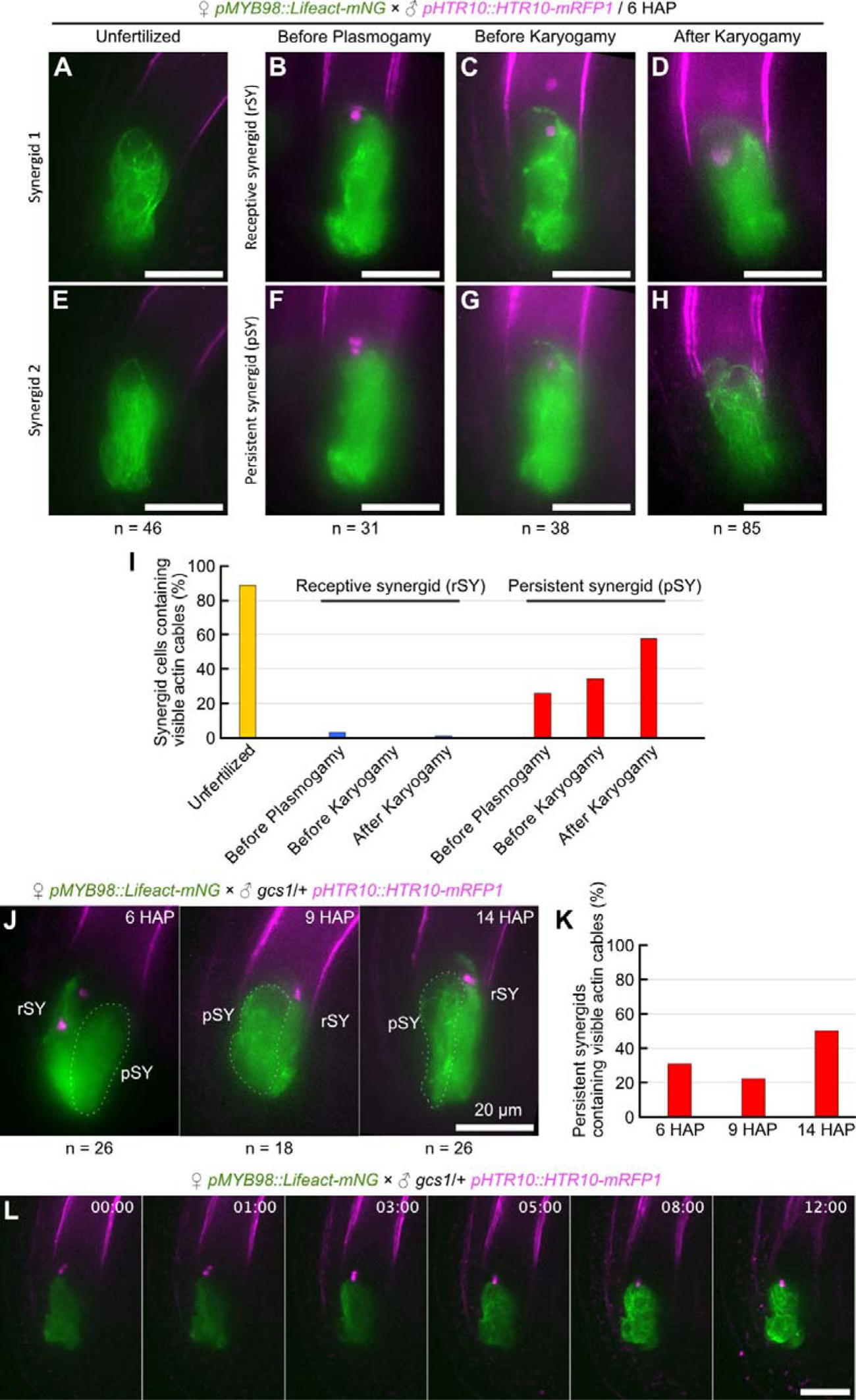
Destruction and reconstruction of F-actin in the persistent synergid. **A–I,** F-actin pattern in the synergid cells during fertilization. Pistils from the *pMYB98::Lifeact-mNG* F-actin marker line were pollinated with pollen from the *pHTR10::HTR10-mRFP* sperm nuclear marker line and the ovules were analyzed at six hours after pollination (6 HAP) by spinning disk confocal microscopy. Images were maximum intensity projections of synergid cells on different focal planes in each Z-series of ovule at unfertilized stage (A, E), before plasmogamy stage (B, F), before karyogamy stage (C, G), and after karyogamy stage (D, H). After pollen tube reception, receptive synergids (rSY) (B, C, D) can be distinguished from persistent synergids (pSY) (E, F, H) by Lifeact-mNG leakage between the egg cell and central cell. The graph shows the percentage of synergid cells containing visible actin cables (I). **J–L,** F-actin pattern of the persistent synergids in the ovules after reception of fertilization-defective *gcs1* sperm cells. Pistils from the *pMYB98::Lifeact-mNG* were pollinated with pollen from a *gcs1* heterozygous plant carrying the *pHTR10::HTR10-mRFP* marker. Confocal images of ovules before plasmogamy at 6, 9, and 14 hours after pollination (J), and percentages of persistent synergids containing visible actin cables (K). Time-lapse imaging of an ovule dissected from a pistil at 6 hours after pollination (L). Scale bars: 20 μm.

In contrast to the receptive synergid, the persistent synergid did not show actin corona-like structures and the actin cables disappeared before plasmogamy (Figure 8F and 8I; n = 31). The percentage of ovules containing visible actin cables slightly increased at the stage before karyogamy (Figure 8G and 8I; 34%, n = 38), and finally reached 58% after karyogamy (Figure 8H and 8I; n = 85). To examine whether actin cable recovery depended on plasmogamy, we used *gcs1*, a T-DNA insertion allele of *GENERATIVE CELL SPECIFIC 1/HAPLESS2* that produces plasmogamy-defective sperm cells (Mori et al., 2006; von Besser et al., 2006). Pistils from *pMYB98::Lifeact-mNG* were pollinated with the pollens from *gcs1* heterozygous plants carrying *pHTR10::HTR10-mRFP1*, and the F-actin pattern in the persistent synergid was analyzed in ovules containing unfused sperm cells at 6, 9, and 14 HAP (Figure 8J and 8K). At 14 HAP, clear actin cables were observed in the ovules more frequently than at earlier HAP time points (Figure 8K), indicating the fertilization-independent nature of actin cable recovery. To elucidate the regeneration process of actin cables, we next performed a time-lapse imaging. Pistils from *pMYB98::Lifeact-mNG* were pollinated with the pollens from *gcs1/+;pHTR10::HTR10-mRFP1*, and 6 HAP ovules cultured in a liquid medium were observed every 15 min by spinning disk confocal microscopy. In fertilized ovules, persistent synergids were absorbed within a few hours and the F-actin pattern was no longer visible due to the rapid dilution of Lifeact-mNeonGreen (Movie 4). In contrast, ovules containing unfused sperm cells maintained the mNeonGreen signal in the persistent synergid through the 12 hours of time-lapse observation (Figure 8L; Movie 5). In these unfertilized ovules, few actin cables emerged ∼3h after observation and the actin cables gradually increased the number. These results indicate that pollen tube discharge induces temporal actin depolymerization and cell-autonomous regeneration of actin cables in the persistent synergid.

## DISCUSSION

The synergid cell has been extensively studied as the key regulator of the initial communication between male and female gametophytes during fertilization in flowering plants. In this study, genetic and pharmacological approaches revealed different roles of microtubules and F-actin in the morphology and functions of synergid cell. Although partial colocalization of microtubules and actin cables was observed in the synergid cell, F-actin disruption did not affect microtubule pattern and *vice versa* (Figure 2; Figure 3; Figure 6; Figure 7; Figure S5). These observations imply independent functions of the two cytoskeletal components in the synergid cell. Indeed, different phenotypes were observed after disruption of microtubule or F-actin formation. Inhibition of microtubule formation by PHS1ΔP expression led to shorter plasma membrane invaginations at the filiform apparatus, whereas the ovules accomplished pollen tube attraction and fertilization normally (Figure 3; Figure 4). On the other hand, F-actin deformation compromised the polarity of synergid cell organelles and impaired the secretion of pollen tube attractants from the filiform apparatus (Figure 6; Figure 7). Therefore, we propose that longitudinal actin cables play a central role in the formation and maintenance of cell polarity, while microtubules have subsidiary roles in filiform apparatus formation.

Our FIB-SEM and confocal microscopy analyses revealed a detailed sponge-like filiform apparatus structure consisting of plasma membranes and cell walls (Figure 1). The porous cell wall pattern was severely impaired by the expression of DN-ACTIN (Figure 6), suggesting that the proper cell wall pattern at the filiform apparatus requires coordination of actin-mediated protein secretion and cell wall formation. The loss of plasma membrane accumulation was also observed in the PHS1ΔP expressing ovules probably due to the stochastic fusions of two synergid cells and central cell (Figure 3). Interestingly, a subset of this type of ovules may accomplish pollen tube attraction, double fertilization, and seed formation (Figure 4). Although details of the synergid-central cell fused cell with multifunction are still unclear, our data indicate the capability of pollen tube attraction and reception in the absence of plasma membrane invaginations at the filiform apparatus. If the filiform apparatus is dispensable from the synergid cell, why this complex structure is conserved among flowering plants? The role of complexity at the filiform apparatus may not be confined to only surface expansion of the protein secretion area. Perhaps, the porous cell walls may serve as a genuine sponge for the temporary storage of signaling molecules to ensure efficient reproduction. For example, this putative signal storage might support accurate pollen tube attraction by buffering fluctuations of protein secretion. Alternatively, it may modulate pollen tube behavior through the release of a large amount of signaling molecules when the growing pollen tube tip pushes the filiform apparatus before pollen tube discharge (Higashiyama et al., 1998; Leshem et al., 2013; Ngo et al., 2014).

Not only pollen tube attractant peptides, but also various pollen tube reception factors such as FERONIA, ANJEA, HERK1, LORELEI, and NORTIA are localized at the plasma membrane of the filiform apparatus (Escobar-Restrepo et al., 2007; Galindo-Trigo et al., 2020; Capron et al., 2008; Tsukamoto et al., 2010; Kessler et al., 2010). NORTIA is relocated from a Golgi-associated compartment to the filiform apparatus during pollen tube reception, implying cargo-dependent spatiotemporal control of polarized protein secretion in the synergid cell. Polarized protein secretion has been extensively studied in tip-growing model cells including the pollen tube, root hair cell, protonema in mosses, and filamentous fungi (Stephan, 2017; Bibeau et al., 2018). In these cells, F-actin is often concentrated beneath the tip region forming the actin fringe, which facilitates the accumulation of secretory vesicles and subsequent control of polarized secretions crucial for tip-growth (Stephan, 2017). Although F-actin concentration was not obvious around the filiform apparatus, their active rearrangement may be involved in polarized protein secretion as actin fringes in tip-growing cells.

The movement toward the filiform apparatus was another feature of F-actin dynamics observed in the synergid cell. This is considerably different from actin dynamics in the egg cell and central cell, where actin cables emerge from the cell periphery and move toward the nucleus (Kawashima et al., 2014; Ohnishi et al., 2014; Ohnishi and Okamoto, 2015). The nucleopetal movement in the female gametes is regulated by an Arp2/3-independent WAVE/SCAR actin nucleation pathway activated by ROP small GTPase and is important for the migration of sperm nuclei after plasmogamy (Ali et al., 2020). Although the female gametophyte cells are originated from the same cell lineage, their functions, morphology, and gene expression are quite different at maturity. Therefore, the synergid cell may present its actin dynamics controlled by a different set of actin modifiers including ROP, Arp2/3, WAVE/SCAR, Formins, and other actin-binding proteins. The F-actin regulator, SCAR2 controls F-actin meshwork movement in the central cell and possibly in the egg cell (Ali et al., 2020). In our previous study, the synergid cells in *myb98* mutant, lacking extensive invaginations characteristic of the filiform apparatus, have partially egg cell-like gene expression and nuclear position. In the *myb98* synergid cell, slight upregulation in *SCAR2* expression (Susaki et al., 2021) supports the hypothesis of cell-specific regulation of actin in the female gametophyte cells.

In our observation, F-actin pattern in the synergid cell changes drastically during fertilization. After pollen tube discharge, fine actin cables disappeared in the ruptured receptive synergid and the synergid-derived Lifeact-mNeonGreen signal appeared as an actin corona (Figure 8). Actin corona was discovered by a Rhodamine-phalloidin staining to observe actin cables around discharged sperm cells prior to double fertilization in *Nicotiana tabacum* (Russell, 1993; Huang and Russell, 1994). Due to its characteristic spatiotemporal pattern, actin corona was speculated to transport the released sperm cells to the area where double fertilization takes place. However, live-imaging of double fertilization in direct delivery of sperm cells within a few seconds, implying actin-independent sperm cell delivery mechanism in *A. thaliana* (Hamamura et al., 2011). In this study, we found the actin corona-like signal accumulation of the Lifeact-mNeonGreen exhibited moderate colocalization with cytosolic tagRFP released from the pollen tube (Figure S6). Actin corona may be an ephemeral cytosol accumulation irrelevant to sperm cell migration.

The persistent synergid displayed temporal disappearance of actin cables after pollen tube discharge, which may pause secretion of pollen tube attractant peptides and be involved in the mechanisms controlling polytubey block (Maruyama and Higashiyama, 2016). Polytubey block is completed by the degeneration of the persistent synergid triggered by double fertilization. The degeneration processes include a cell-fusion between the persistent synergid and endosperm (Maruyama et al., 2015) and degeneration of the persistent synergid nucleus, which is mediated by the EIN3 and EIL1 transcription factors (Völz et al., 2013; Li et al., 2022; Heydlauff et al., 2022) and FIS-class polycomb repressive complex (Maruyama et al., 2013; Maruyama et al., 2015). Compared to these late degeneration processes, the disappearance of actin cables may be an earlier and fertilization-independent alteration that temporarily inactivates persistent synergid functions (Figure 8F, 8I, 8J–K). Recent reports have demonstrated several early polytubey block pathways that diminish the attractant signals around the micropyle. Specifically, FERONIA-dependent NO production accompanying with a deposition of de-esterified pectin inactivates AtLURE1 peptides via post-translational nitrosation upon pollen tube arrival (Duan et al., 2020). Active AtLURE1 peptides are further reduced by ECS1 and ECS2, which are egg cell-specific aspartic endopeptidases secreted immediately after plasmogamy (Yu et al., 2021). Together with these attractant peptides elimination systems, inhibition of protein secretion in the persistent synergid should play a key role in the early polytubey block. Termination of AtLURE1 secretion and intracellular accumulation of AtLURE1 in the synergid cell are also induced by elevation of NO signaling during pollen tube arrival (Duan et al., 2020). Although the precise mechanism is still unknown, the NO-mediated termination of AtLURE1 secretion may be controlled by the temporal destruction of actin cables. Further studies using inhibitors and mutants affecting NO production and F-actin formation are required to confirm this hypothesis.

Interestingly, the persistent synergid displayed actin cables regeneration, which was more evident in unfertilized ovules after reception of the plasmogamy-defective *gcs1* mutant pollen tube (Figure 8L). When the first pollen tube fails to fertilize the female gametes, the ovules stop the degeneration of the persistent synergid and begin to attract a second pollen tube within ∼5h (Kasahara et al., 2012; Duan et al., 2020; Zhong et al., 2022). The time window between the arrival of the first and second pollen tubes coincides with the period of actin cable disappearance in the persistent synergid, suggesting that actin cables dynamics of destruction-reconstruction function as a courtship-pause timer allowing the persistent synergid to attract the second pollen tube. Occasionally, polytubey causes polyspermy, which might provide the offspring agronomically useful traits inherited from three parents (Nakel et al., 2017; Grossniklaus, 2017; Mao et al, 2020). To increase the efficiency of such polyspermy breeding, cell biology of persistent synergid reactivation would become important in the future. Our discovery of the actin cable regeneration could be the first step to understanding molecular mechanism of the persistent synergid reactivation.

## Methods

### Plant materials and growth conditions

Columbia-0 (Col-0) was used as the wild-type strain. The *gcs1* (SALK_135694) heterozygous mutant that was also homozygous for the *pHTR10::HTR10-mRFP* has been described previously (Kasahara et al, 2012). Seeds were surface sterilized with sterilization solution (2% PLANT PRESERVATIVE MIXTURE ^TM^, 0.1% Tween 20) and incubated at 4°C for several days and sown on Murashige and Skoog (MS) medium containing 1% sucrose and appropriate antibiotics. Approximately 10 days after germination, plants were transferred to soil. Plants were grown at 22°C under continuous light or standard long day conditions.

### Plasmids and transgenic plants

The *pRPS5A::H2B-tdTomato* plasmid was kindly provided by D. Kurihara (Nagoya Univ.) (Maruyama et al., 2013). pDM286, a pGWB501 destination vector (Nakagawa et al., 2007) containing the *MYB98* promoter, has been reported previously (Maruyama et al., 2015). The *pAGL80::tagRFP-TUA5*, pDM230 (a binary vector carrying the *pLAT52::tagRFP*) and the pSNA128 (a pGWB501 destination vector containing 1,062 bp upstream of the coding region of the *ES2* (At1g26795) (Pagnussat et al, 2007; Hwang et al., 2019)) were gifts from Shuh-ichi Nishikawa (Niigata Univ.). The primers used for plasmid constructions are listed in Table S1.

The *PHS1ΔP*, a mutant gene of C-terminus-truncated PROPYZAMIDE-HYPERSENSITIVE 1 (85**–**700 a.a.) (Fujita et al., 2013), was amplified from *Arabidopsis* seedling cDNA by a PCR using primers ‘PHS1dP_F+SalI-TOPO’ and ‘PHS1dP_R+AscI-stop’, and introduced into pENTR/D-TOPO vector (Invitrogen) to generate the HTv1243. An LR recombination between the pDM286 and HTv1243 produced pDM580, a binary vector containing the *pMYB98::PHS1ΔP*.

The HTv1313 and HTv1316, two pollen tube attractant reporter plasmids were cloned as follows. The coding sequence of yellow fluorescent protein Citrine was introduced into pPZP211 vector by replacing mRuby2 of pPZP211Ru (Takeuchi and Higashiyama, 2016), resulting in pPZP211Cit. The entire genomic sequence of *AtLURE1.2* including the protein-coding and upstream promoter sequences was amplified from *Arabidopsis* genomic DNA by a PCR using primers ‘AtLURE1.2_F+SalIGA’ and ‘AtLURE1.2_-stopR+AscIGA’, and introduced into the SalI/AscI site of the pPZP211Cit to produce the HTv1313, a binary vector harboring the *pAtLURE1.2::AtLURE1.2-Citrine*. Similarly, entire genomic sequence of the CRP810_2.3/XIUIQIU4/TIC3 was amplified by a PCR using primers ‘810_2.3_F+SalIGA’ and ‘810_2.3_-stopR+AscIGA’, and introduced into the SalI/AscI site of the pPZP211Cit to produce the HTv1316, a binary vector harboring the *XIUQIU4::XIUQIU4-Citrine*.

To generate the synergid cell-specific F-actin reporter genes, a DNA fragment of *mNeonGreen*- or *mRUBY3*-tagged *Lifeact* was amplified by PCR using primers ‘pENTR LifeActFP_F’ and ‘FP_R’ and cloned into pENTR/D-TOPO to produce pOR099 or pOR102. An LR recombination between the pDM286 and pOR099 produced pDM522, a binary vector containing the *pMYB98::Lifeact-mNG*, and an LR recombination between the pDM286 and pOR102 produced pDM523, a binary vector containing the *pMYB98::Lifeact-mRUBY3*.

To generate the synergid cell-specific microtubule reporter gene, a DNA fragment of *mTFP1* was amplified by PCR using primers ‘pENTR_CFP_F’ and ‘FP_PstI_R’ and cloned into pENTR/D-TOPO. Then, *TUA5* sequence in the *pAGL80::tagRFP-TUA5* was introduced into *Nco*I/*Pst*I site of the *mTFP1-*containing entry vector to produce pOR103. The *Citrine* coding sequence was amplified from the HTv1313 by a PCR using primers ‘pENTR_cacc_XFP_F’ and ‘FP_PstI_R’, and substituted with *mTFP1* in the pOR103 by *Nco*I/*Pst*I digestion and subsequent ligation to generate pOR177. An LR recombination between the pDM286 and pOR177 produced pDM680, a binary vector containing the *pMYB98::Citrine-TUA5*.

Plasma membrane markers were constructed as follows. The pOR084, an entry clone containing *mRUBY2-SYP132*, was constructed using NEBuilder HiFi DNA Assembly Cloning Kit (New England Biolabs, MA, USA) by multi-DNA fragment assembly between (i) *mRUBY2*-containing pENTR/D-TOPO (Thermo Fisher Scientific, MA, USA) plasmid backbone amplified by a primer set ‘FP_SpeI_F’ and ‘FP_EcoRI_R’ and (ii) *SYP132* (At5g08080) cDNA sequence amplified by a primer set ‘SYP132_EcoRI_F’ and ‘SYP132_SpeI_R’. pENTR/D-TOPO plasmid backbone with the *SYP132* was amplified from the pOR084 by a PCR using primers ‘GFP_1-18_R’ and ‘XFP_C-term_F’. Resulting linearized plasmid DNA was mixed with three mNeonGreen DNA fragments independently amplified by the following three primer sets, ‘GFP F’ and ‘Linker1_XFP_R’, ‘Linker1_XFP_F’ and ‘Linker2_XFP_R’, or ‘Linker2_XFP_F’ and ‘FP_R’, and multi-DNA fragment assembly by the NEBuilder HiFi DNA Assembly Cloning Kit was performed to produce pOR129. An LR recombination of the pSNA128 and pOR129 generated *pES2::3×mNG-SYP132*. An LR recombination of the pDM286 and pOR129 generated *pMYB98::3×mNG-SYP132*. Two plasmids containing wild-type *ACTIN8* (At1g49240) or dominant negative mutant of *ACTIN8* (Kato et al., 2010) downstream of the *MYB98* promoter, were constructed by LR recombination between the pDM286 and the entry clones carrying wild-type or dominant negative *ACTIN8* that described in previously (Kawashima et al., 2014).

Agrobacterium-mediated plant transformation was performed by the floral dipping method using Agrobacterium strain GV3101 (Clough and Bent, 1998). Transformants were selected on MS medium containing 50 μg/mL hygromycin B or 50 μg/mL kanamycin.

### Analysis of mature ovules

Pistils were harvested one to two days after emasculation and the carpel walls were removed on a glass slide. Then, the pistils were split into half and mounted on 5% sucrose or an ovule culture medium containing Nitsch basal salt mixture (Duchefa) and 5% trehalose dihydrate (Wako Fujifilm) (Gooh et al., 2015). In pharmacological assay, Latrunculin A or Oryzalin were dissolved in DMSO to make 2.5 mM stock solutions and stored at −20°C. These stock solutions were thawed and diluted 250 times by the ovule culture media before use (final concentration, 100 μM). Mature ovules were incubated in these inhibitor-containing media or 0.4% DMSO (negative control), and confocal images were obtained by Leica SP8 TCS (Leica, Wetzlar, Germany). For time-lapse imaging, ovules were dissected from pistils in a droplet of ovule culture medium containing Nitsch basal salt mixture (Duchefa) and 5% trehalose dihydrate (Wako Fujifilm) (Gooh et al., 2015), and transferred to a glass bottom dish (D141410; Matsunami Glass IND., LTD., Japan). XY positions of the ovules were recorded by bright field observation, and time-lapse confocal images were captured using IX73 inverted microscope (Olympus, Tokyo, Japan) equipped with a spinning disk confocal scanning unit (CSU-W1; Yokogawa, Tokyo, Japan) and an sCMOS camera (Zyla 4.2; Andor, Belfast, Northern Ireland).

### Analysis of seed phenotypes

Siliques at eight day-after-pollination were attached on a glass slide by double-sided tape and the upper side of the carpel walls were removed by an ophthalmic knife. The samples were imaged by a dissecting microscope Leica E4 W (Leica, Wetzlar, Germany). To investigate frequency of normal seeds, we harvested fully matured siliques and counted normal seeds and undeveloped seeds under dissecting microscope.

### Histochemistry

CongoRed staining was performed as previously described by Kasahara et al., (2005) with slight modification. Dissected pistils at 1 day-after-pollination were treated with 0.4% CongoRed Solution for 1 min on a glass slide and washed several times with 5% sucrose. Immuno-Staining of AtLURE1.2 was performed as reported previously (Takeuchi and Higashiyama, 2012). These samples were observed by an upright microscope (Axio Imager. M1; Zeiss, Jera, Germany) using filter sets for DsRed or GFP. Epi-fluorescent images of those pistils were captured by the IX73 inverted microscope (Olympus, Tokyo, Japan).

### FIB-SEM

FIB-SEM sample blocks were prepared as follows. Wild-type Col-0 ovules were fixed overnight in a fixation solution (4% paraformaldehyde, 2% glutaraldehyde, 0.05 M cacodylate buffer), then washed three times with 0.05 M cacodylate buffer. The samples were subjected to postfixation with osmium tetroxide in 0.05 M cacodylate buffer at 4°C for 1h and washed five times with distilled water. After incubation with 1% thiocarbohydrazide for 20 min and five rounds of wash with distilled water, the ovules were treated with 2% aqueous osmium tetroxide at room temperature for 1h and washed three times with distilled water. The samples were stained overnight with 1% aqueous uranyl acetate at 4°C, then treated with 0.03M lead aspartate at 60°C for 30 min. The tissue was then dehydrated with increasing ethanol concentrations (50%, 70%, 90%, 100%), transferred into propylene oxide, infiltrated, and embedded in Quetol 651. Specimens were analyzed by FIB-SEM as described previously (Oi et al., 2017). Briefly, the surfaces of the embedded ovules were exposed using a diamond knife on an ultramicrotome (EM UC6; Leica, Wetzlar, Germany), and the resin blocks were trimmed to a cuboid and were attached on the standard specimen stage for FIB-SEM as the exposed ovules faced the SEM column. The specimens were coated with a thin layer using a carbon coater (CADE-E, Meiwafosis, Tokyo, Japan) to prevent electron charging. The serial electron micrographs were obtained with a FIB-SEM (MI-4000L, Hitachi, Tokyo, Japan). The FIB was operated with following conditions: accelerating voltage, 30 kV; beam current for milling to prepare or to cut, 1.6 or 1.2 nA, respectively; cutting interval (z-step), 25 nm. The SEM was operated with following conditions: accelerating voltage, 1.0 kV; working distance, 2 mm; image size, 2000×2000 pixels; color depth, 8-bit (256 grey scales); pixel size, 25 nm per pixel. Secondary and backscattered electrons were detected by the detector in the SEM column (U+EsB). The images of serial sections were processed using the software ‘Fiji’ (http://fiji.sc/Fiji) (Schindelin et al., 2012); grey scale was inverted, brightness and contrast were adjusted, the images were aligned using the ‘Register Virtual Stack Slices’ (Translation—Translation -no deformation) tool, and then the region of the filiform apparatus was cropped (image size, 400 ×400 pixels). The outline of cell walls of two synergid cells in all 2D images were traced manually using the software ‘PaintTool SAI’ (ver. 1, Systemax, Japan). The inside region of the two synergid cells were binarized to detect the filiform apparatus and to reconstruct it into the 3D model using the software ‘Image-Pro Premier 3D’ (ver. 9.3, Media Cybernetics, USA); the surface rendering model was reconstructed with subsampling (64M voxel) and smoothing (low-pass filter [3:3:3]), and then their surface area were calculated with the function ‘3D Measure’.

### Transmission electron microscopy

Emasculated pistils from the transgenic plant homozygous for the *pMYB98::PHS1ΔP*, *pRPS5A::H2B-tdTomato*, and *pES2::3×mNG-SYP132*, were dissected on glass slides and analyzed by an upright microscope (Axio Imager. M1; Zeiss, Jera, Germany) to select the ovules containing single synergid cell. Similarly, normal and abnormal ovules were selected from the *pAtLURE1.2::AtLURE1.2-Citrine transgenic plant hemizygous for the pMYB98::DN-ACTIN*. The ovule selection procedure was skipped when we prepared wild-type controls of the cytoskeleton-disrupted transgenic plants. The ovules were fixed in a solution containing 4% paraformaldehyde, 2% glutaraldehyde, and 50 mM sodium cacodylate at pH7.4 for several days at 4°C. The samples were washed in buffer and post-fixed for 6 h in 2% aqueous osmium tetroxide at 4°C. The specimens were then dehydrated in a graded ethanol series, transferred into propylene oxide, infiltrated, and embedded in Quetol 651. Series of thin-sections (80 nm) were stained with 2% aqueous uranyl acetate and lead citrate, and examined at 80 kV under a JEOL JEM 1400 Plus electron microscope (JEOL Ltd.). Digital images were taken with a CCD camera (VELETA; Olympus Soft Imaging Solutions).

### Analysis of F-actin flow

Time-lapse images of F-actin dynamics in synergid cell expressing *pMYB98::Lifeact-mNG* were captured by spinning disk confocal microscopy with 30 sec intervals (total 21 frames; 600 sec) and multiple z-planes (total 9 planes, 1 μm intervals). The image was deconvoluted using the constrained iterated deconvolution function of CellSens Dimension Desktop 3.2 (Olympus). Background noise was removed using the subtract background function in Fiji (Schindelin et al., 2012). Three z-planes including the peri-nuclear areas of the time-lapse images were projected and then used for analysis of F-actin flow in the synergid cell. To analyze the F-actin flow, flow velocity vectors (v_x_,v_y_) were obtained from spatio-temporal image correlation spectroscopy analysis using MATLAB (MATLAB Inc.) scripts (Ashdown et al., 2015) with parameter settings: pixelSize 0.259, timeframe 30, tauLimit 4, filtering FourierWhole, MoveAverage 21, ROIsize 16, ROIshift 4, TOIsize 3, TOIshift 1 as described in the manual (Ashdown et al., 2015). After the flow velocity vectors were acquired from each analysis area, all angles of the flow velocity vectors from the entire analyzed frames were calculated and normalized as probability using a custom MATLAB script to get direction distribution of the F-actin flow.

## Conflict of interest

The authors declare that the research was conducted in the absence of any commercial or financial relationships that could be construed as a potential conflict of interest.

## Author contributions

D.M. and T. Ka. conceived this project; D.S. obtained confocal images in Figure S1, 2 and S2; T.O. conducted FIB-SEM analysis and image processing in Figure 1 and Movie 1; J.M.S., T. Ka., and D.S. analyzed F-actin dynamics in Figure 2, S2 and Movie 3. R.I. and D.M. analyzed fertility of DN-ACTIN expressing lines; R.I and H.T. contributed immunostaining in Figure S2; N.S. analyzed the invagination length in Figure1; D.M. obtained other data and directed this project; D.M. and D.S. wrote the manuscript and all authors contributed to edit the manuscript.

## Acknowledgements

FIB-SEM observation was supported by Nagoya University microstructural characterization platform as a program of “Nanotechnology Platform” of the Ministry of Education, Culture, Sports, Science and Technology (MEXT), Japan, and we especially thank S. Arai and S. Enomoto (Nagoya Univ.) for technical support. We acknowledge B. Raj Thapa (Univ. of Kentucky) for supporting on the F-actin movement quantification, M. Tsukatani for assistance in image analysis, H. Ikeda for assistance in preparing materials, D. Kurihara (Nagoya Univ.), S. Nishikawa (Niigata Univ.) and T. Nakagawa (Shimane Univ.) for providing plasmids. We would like to thank Editage (www.editage.com) for English language editing.

## Funding

This work was supported by TOYOAKI Scholarship Foundation, Japan Society for the Promotion of Science (JSPS) KAKENHI [Grant Numbers: 17H05846, 19H04869, 20H03280, 20H05778 and 20H05781 to D.M.; 19K16172 and 22K15145 to D.S.; 18K14729 and 20K15817 to H.T.], by the Grant for academic research from Yokohama City University (to D.M.), by the grant for 2016–2022 Research Development Fund of Yokohama City University (to D.M.), and by National Science Foundation Grant IOS-1928836 (to T.Ka.).

---

**Figure S1.**
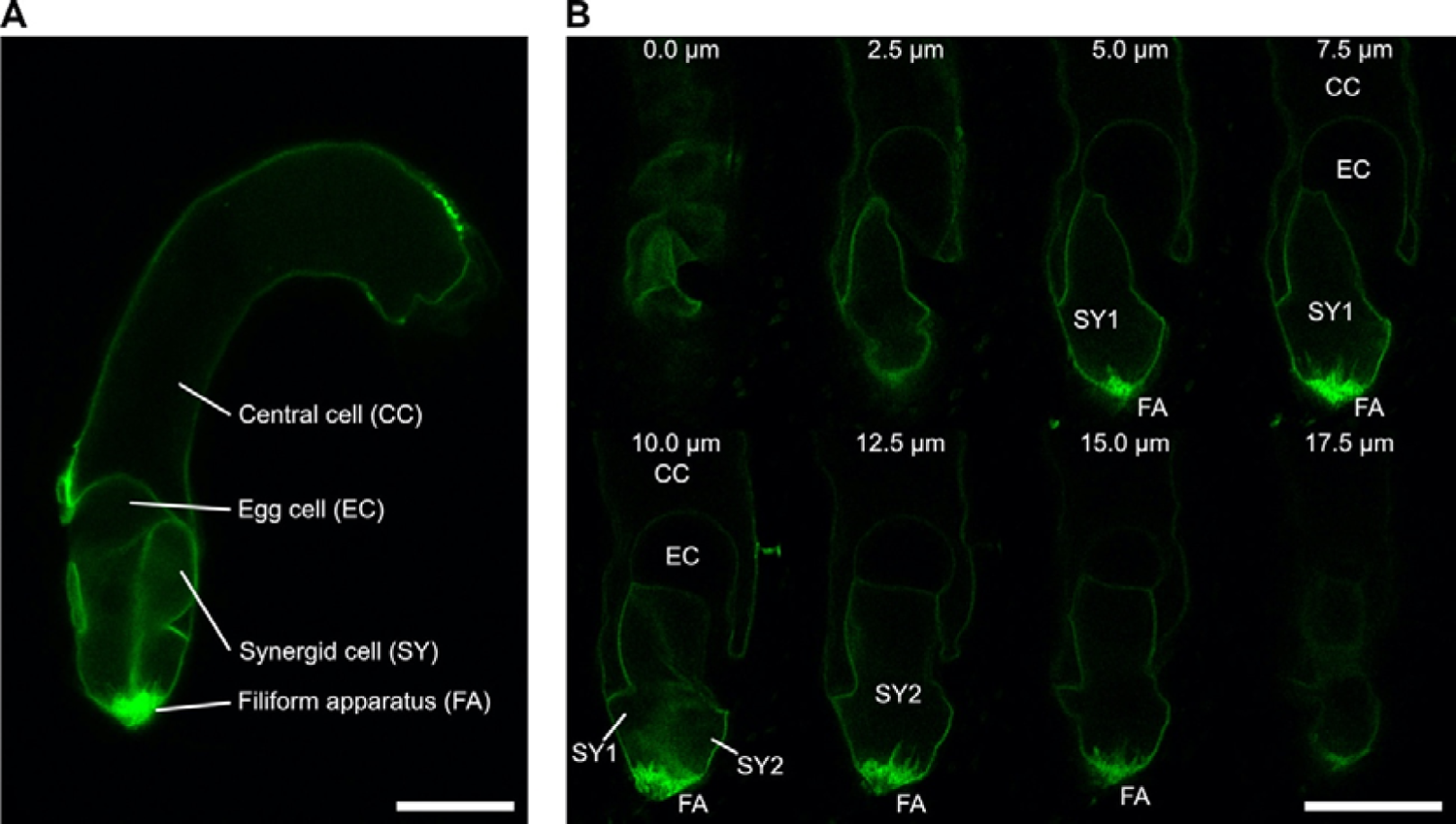
Confocal images of mature ovules from the *pES2::3×mNG-SYP132* (*ENS*) plasma membrane marker line. **A,** An image of center plane. **B,** Z-series images of a single ovule. Scale bars: 20 μm.

**Figure S2.**
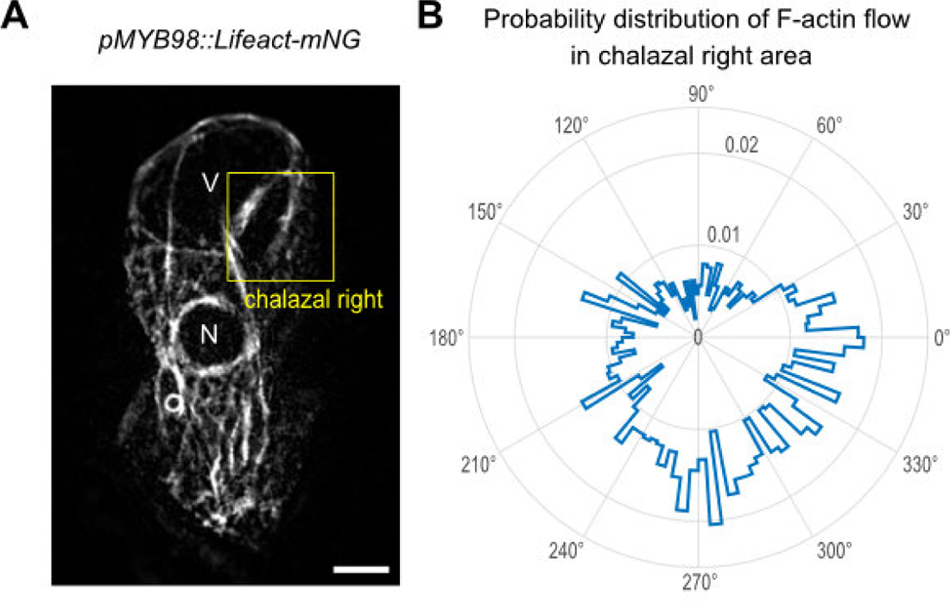
F-actin flow in the chalazal right area of synergid cell. **A,** The first frame of time-lapse imaging of F-actin (Grey) dynamics in the synergid cell expressing *pMYB98::Lifeact-mNG* (Figure 2B). The yellow box indicates the chalazal right area where F-actin flow was analyzed as shown in **B**. V, vacuole; N, nucleus; Scale bar, 10 μm. **B,** Radar plot displaying the probability distribution of F-actin flow directions in the chalazal right area (the yellow box in **A**). The directions are calculated from the angles of F-actin flow vectors obtained from spatiotemporal image correlation spectroscopy. The probability is calculated from cumulative frequency of the angles in F-actin flow vectors from the entire frames of time-lapse imaging and scaled in the radial axes on the radar plot.

**Figure S3.**
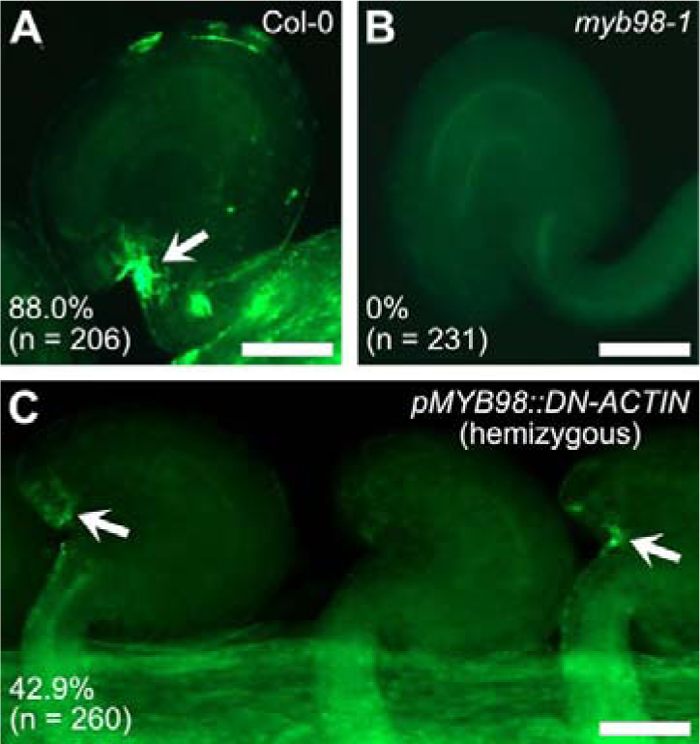
Immunostaining of AtLURE1.2. **A,** Wild-type ovule. **B,** *myb98-1* mutant ovule. **C,** Ovules in the *pMYB98::DN-ACTIN* hemizygous plant. Arrows represents AtLURE1.2 accumulation at the micropyle. Percentages of the AtLURE1.2 accumulation were shown in the bottom-left with the number of total ovules analyzed. Scale bars: 50 μm.

**Figure S4.**
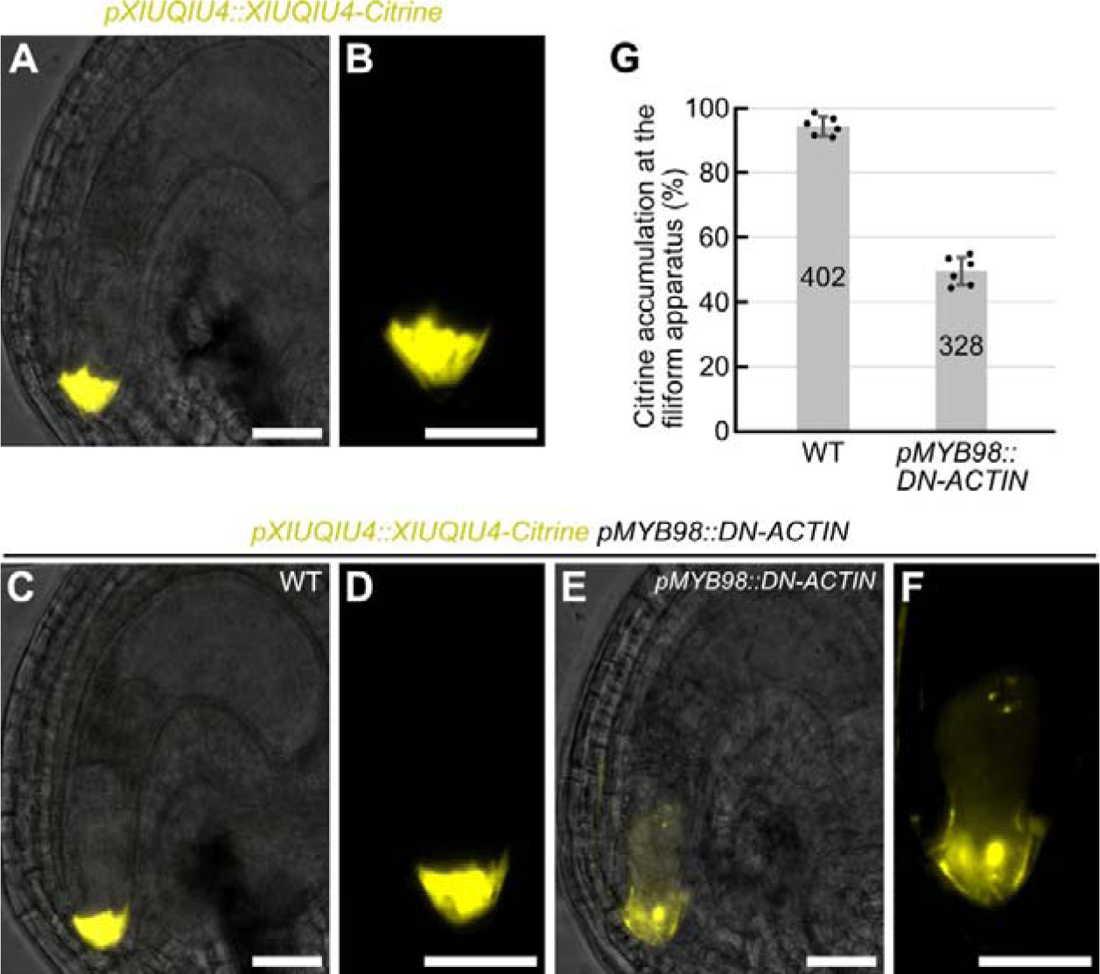
F-actin is necessary for directional XIUQIU4 secretion. **A–F,** Fluorescent signals of *pXIUQIU4::XIUQIU4-Citrine* pollen tube attractant marker in wild-type plant (A, B), and *pMYB98::DN-ACTIN* hemizygous plant (line 1) (C–F). Merges of fluorescent images and differential interference contrast images were shown in (A), (C), (E). Magnified synergid cells were shown in (B), (D), (F). The *pMYB98::DN-ACTIN* hemizygous plant showed segregation of wild-type ovules with normal fluorescent pattern (C, D), and abnormal ovules with little XIUQIU4-Citrine signal at the filiform apparatus (E, F). **G,** Frequency of ovules displaying normal XIUQIU4-Citrine accumulation at the filiform apparatus analyzed in (A) to (F). Scale bars: 20 μm.

**Figure S5.**
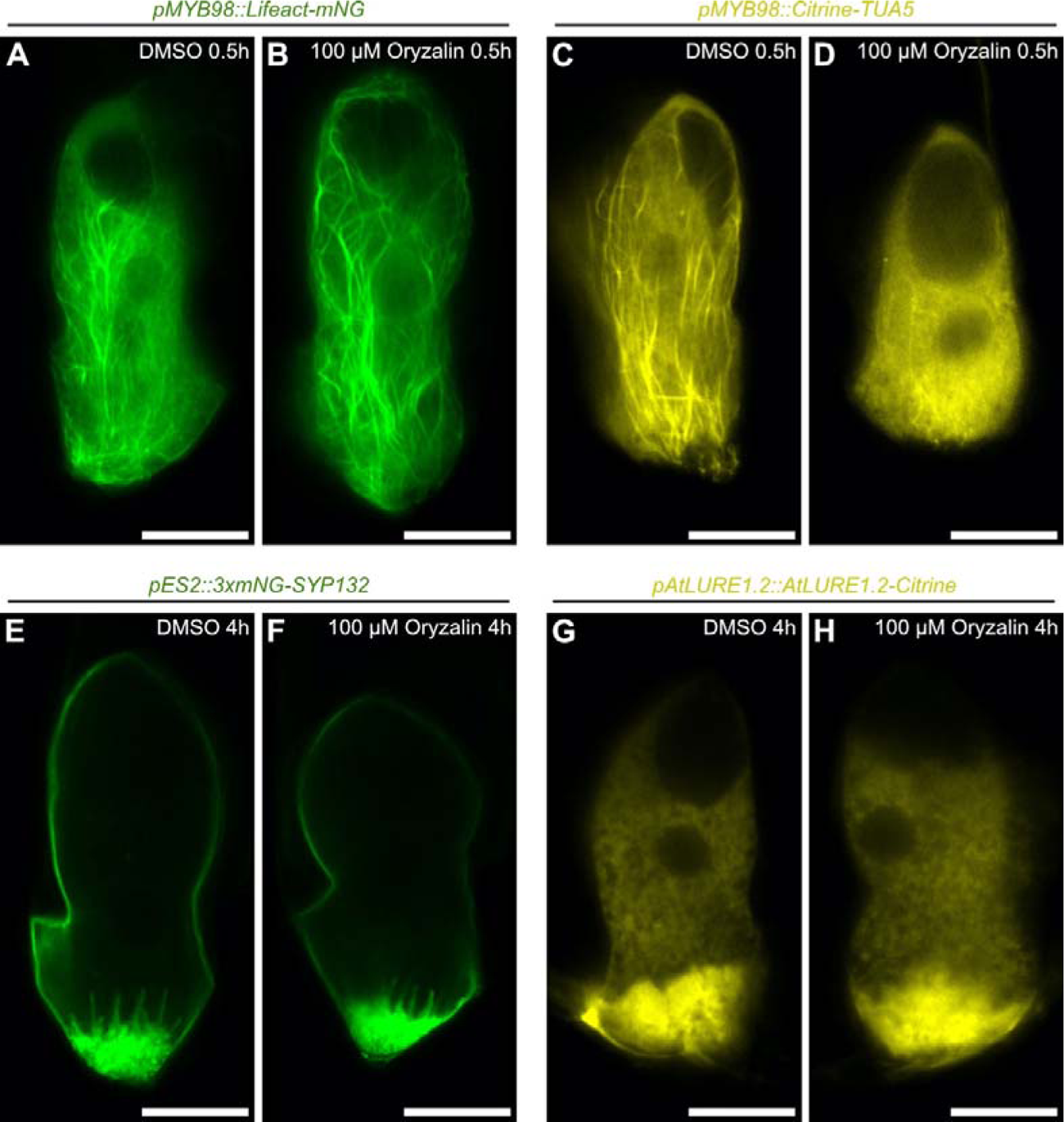
Transient microtubule depolymerization. **A–H,** Confocal images of synergid cells in the *pMYB98::Lifeact-mNG* (F-actin marker line) (A, B), *pMYB98::Citrine-TUA5* (Microtubule marker line) (C, D), *pES2::3×mNG-SYP132* (plasma membrane marker line) (E, F), and *pAtLURE1.2::AtLURE1.2-Citrine* (pollen tube attractant marker line) (G, H), cultured in a control medium containing 0.4% DMSO (A, C, E, G), or a medium containing 0.4% DMSO and 100 μM Oryzalin (B, D, F, H). Duration of the ovule culture is shown in the top-right. Scale bars: 20 μm.

**Figure S6.**
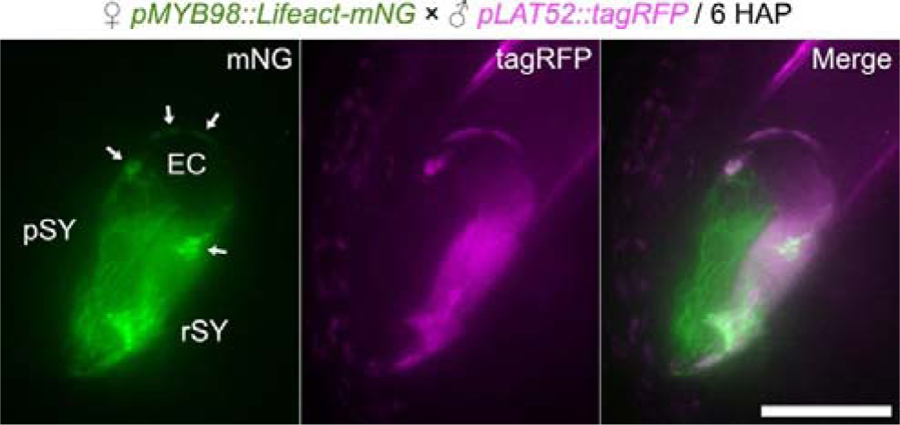
Moderate colocalization of actin corona-like structure and pollen tube cytosol. Pistils from the *pMYB98::Lifeact-mNG* F-actin marker line were pollinated with pollen from the *pLAT52::tagRFP* pollen tube cytosolic marker line, and confocal images of an ovule were captured at six hour after pollination. Arrows represents the corona-like structures labeled with Lifeact-mNeonGreen. Abbreviations: EC, egg cell; rSY, receptive synergid; pSy, egg cell. Scale bar: 20 μm.

**Table S1.**
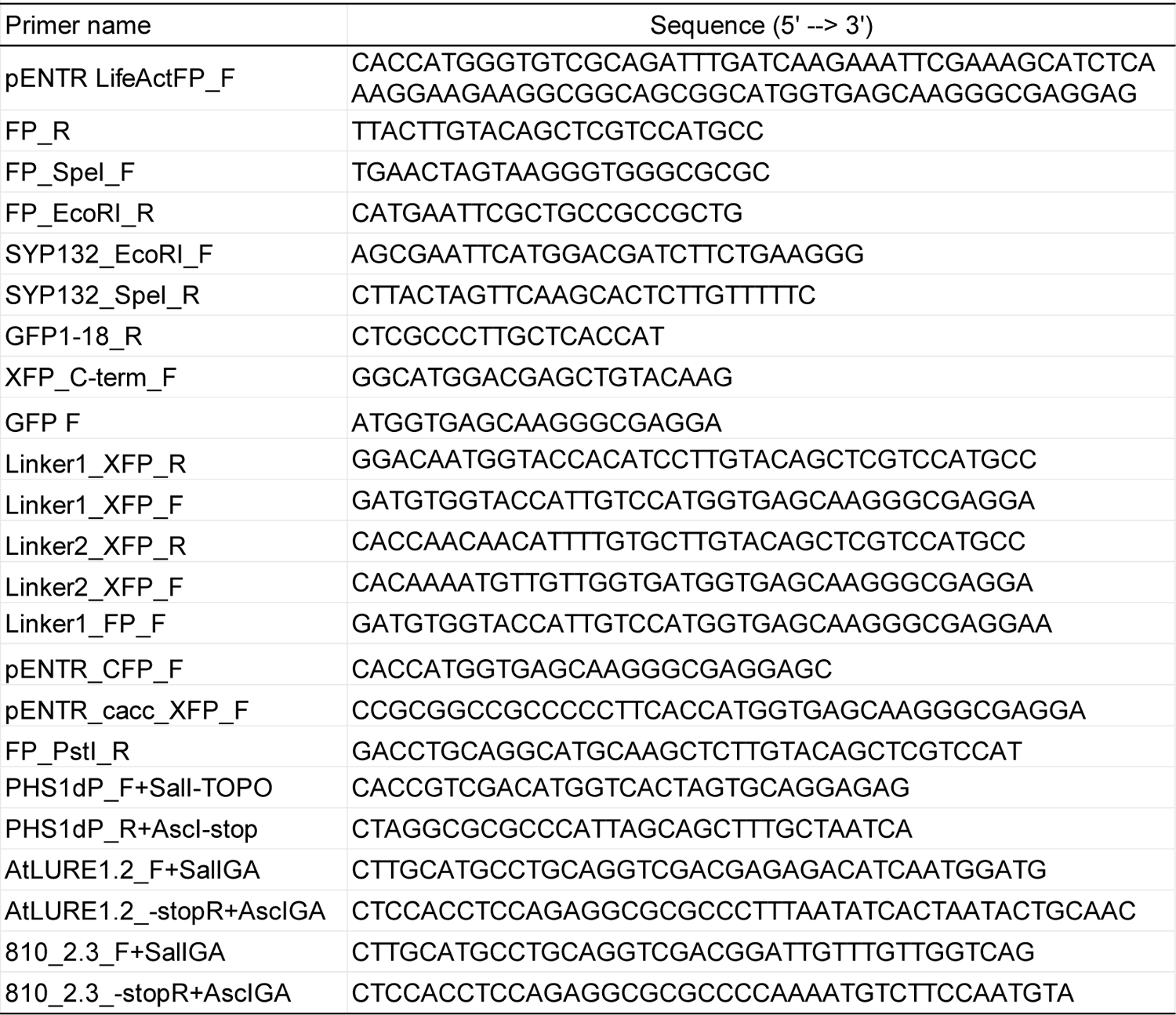
Primers used in this study.

## Captions for Movies

**Movie 1** A 3D image of cell walls at the filiform apparatus reconstructed from serial sections obtained by FIB-SEM.

**Movie 2** A confocal z-series images of filiform apparatus in the *pMYB98::3×mNG-SYP132* plasma membrane marker line.

**Movie 3** Dynamics of F-actin in a synergid cell of mature ovule. Ovules from the *pMYB98::Lifeact-mNG* F-actin marker line were incubated in a liquid culture medium and subjected to time-lapse imaging using spinning disk confocal microscopy.

**Movie 4** F-actin dynamics of persistent synergids in two fertilized ovules. Pistils from the *pMYB98::Lifeact-mNG* were pollinated with pollen from a *gcs1* heterozygous plant carrying the *pHTR10::HTR10-mRFP* marker, and fertilized ovules displaying dispersed RFP signals of decondensed sperm-derived chromosomes in zygote nucleus and endosperm nucleus were analyzed by time-lapse imaging.

**Movie 5** F-actin dynamics of persistent synergids in three unfertilized ovules. Pistils from the *pMYB98::Lifeact-mNG* were pollinated with pollen from a *gcs1* heterozygous plant carrying the *pHTR10::HTR10-mRFP* marker, and unfertilized ovules containing a pair of punctate RFP signals of unfused sperm nuclei were analyzed by time-lapse imaging.

## Notes

### Competing Interest Statement

The authors have declared no competing interest.

